# Extensive and diverse patterns of cell death sculpt neural networks in insects

**DOI:** 10.1101/626465

**Authors:** Sinziana Pop, Chin-Lin Chen, Connor J Sproston, Shu Kondo, Pavan Ramdya, Darren W Williams

## Abstract

Changes to the structure and function of neural networks are thought to underlie the evolutionary adaptation of animal behaviours. Among the many developmental phenomena that generate change programmed cell death appears to play a key role. We show that cell death occurs continuously throughout insect neurogenesis and happens soon after neurons are born. Focusing on two dipterans which have lost flight during evolution we reveal that reductions in populations of flight interneurons are likely caused by increased cell death during development.

Mimicking an evolutionary role for increasing cell numbers, we artificially block programmed cell death in the medial neuroblast lineage in *Drosophila melanogaster*, which results in the production of ‘undead’ neurons with complex arborisations and distinct neurotransmitter identities. Activation of these ‘undead’ neurons and recordings of neural activity in behaving animals demonstrate that they are functional. Our findings suggest that the evolutionary modulation of death-based patterning could generate novel network configurations.

## INTRODUCTION

Nervous systems are exquisitely adapted to the biomechanical and ecological environments in which they operate. How they evolve to be this way is largely unknown. Such changes can occur through modifications in receptor tuning, transmitter/receptor repertoires, neuronal excitability, neuromodulation, structural connectivity, or in the number of neurons within specific regions of the central nervous system (CNS). The differences seen in networks, over an evolutionary timescale, ultimately result from heritable changes in developmental processes (Horder, 1989). Advancing our knowledge of the mechanisms of neural development using comparative approaches will help us understand how specific elements can be modified, how new ‘circuits’ and behaviours evolve, and will ultimately lead to a better understanding of how nervous systems function (Ramdya and Benton, 2010). Studies comparing the nervous systems of mammalian species that occupy diverse ecological niches reveal clear differences in the number of cells within homologous brain regions (Herculano-Houzel et al., 2014). Such differences have occurred either through expansion or reduction of specific cell populations, through changes in proliferation or apoptotic programmed cell death (PCD) during development (Charvet et al., 2011). Most studies of nervous system evolution have focused on stem cell identity and the role of differential proliferation dynamics (Truman and Ball, 1998; Rakic, 2009; Biffar and Stollewerk, 2014). While one recent study has elegantly shown a role for PCD in the evolution of peripheral olfactory sensory neurons in drosophilids and mosquitoes (Prieto-Godino et al., 2020), how changes in cell death can modify central circuits still remains an open question.

In insects the number and arrangement of neural progenitor cells that generate central neurons (termed neuroblasts, NBs) is highly conserved despite a remarkable diversity of insect body plans and behaviors (Wheeler, 1891; Bate, 1976; Hartenstein and Campos-Ortega, 1984; Doe and Goodman, 1985; Tamarelle et al., 1985; Booker and Truman, 1987; Truman and Bate, 1988; Shepherd and Bate, 1990; Doe, 1992; Truman, 1996; Truman and Ball, 1998; Biffar and Stollewerk, 2014). In the ventral nerve cord (VNC – functionally equivalent to the vertebrate spinal cord), the medial neuroblast (MNB) in particular stands out as an unpaired NB located at the midline in the posterior end of each segment, while all other NBs come in pairs arranged in a bilaterally symmetric array across the midline (**Figure 1A,B,C**). Its easy-to-locate position made the MNB identifiable in all developing insects described from as early as 1891 by W.M. Wheeler (Wheeler, 1891), and spanning all insect orders from wingless silverfish to locusts, beetles, moths, and flies (Bate, 1976; Hartenstein and Campos-Ortega, 1984; Doe and Goodman, 1985; Tamarelle et al., 1985; Booker and Truman, 1987; Truman and Bate, 1988; Shepherd and Bate, 1990; Doe, 1992; Truman and Ball, 1998; Biffar and Stollewerk, 2014). The MNB gives rise to two populations of neurons, one GABAergic and one octopaminergic, which are also homologous across insects (Rowell, 1976; Siegler et al., 1991, 2001; Thompson and Siegler, 1991; Campbell et al., 1995; Stevenson and Spörhase-Eichmann, 1995; Siegler and Pankhaniya, 1997; Witten and Truman, 1998; Jia and Siegler, 2002; Pflüger and Stevenson, 2005; Lacin et al., 2019). There appears to be a relationship between cell number and function in these populations. Flying insects have greater numbers of octopaminergic neurons within segments that control wings (**Figure 1D**; Stevenson and Spörhase-Eichmann, 1995), while grasshoppers have more GABAergic neurons in the fused metathoracic/abdominal ganglia, where they receive auditory input from the abdomen (**Figure 1D**; Witten and Truman, 1998; Thompson and Siegler 1991). Alongside differences in numbers of the same cell type between segments and species, numbers of GABAergic and octopaminergic neurons found in one segment are never equal (**Figure 1D,E**). This is especially intriguing as during development each GABAergic neuron is a sister cell to an octopaminergic neuron, arising from one cell division and are produced in equal number. (**Figure 1F**). The greater number of GABAergic cells in each segment results from PCD targeting octopaminergic neurons in both grasshoppers (Jia and Siegler, 2002) and fruit flies (Truman et al., 2010). The repeated elimination of one neuronal sibling over the other is not restricted to the MNB lineage.

**Figure 1.**
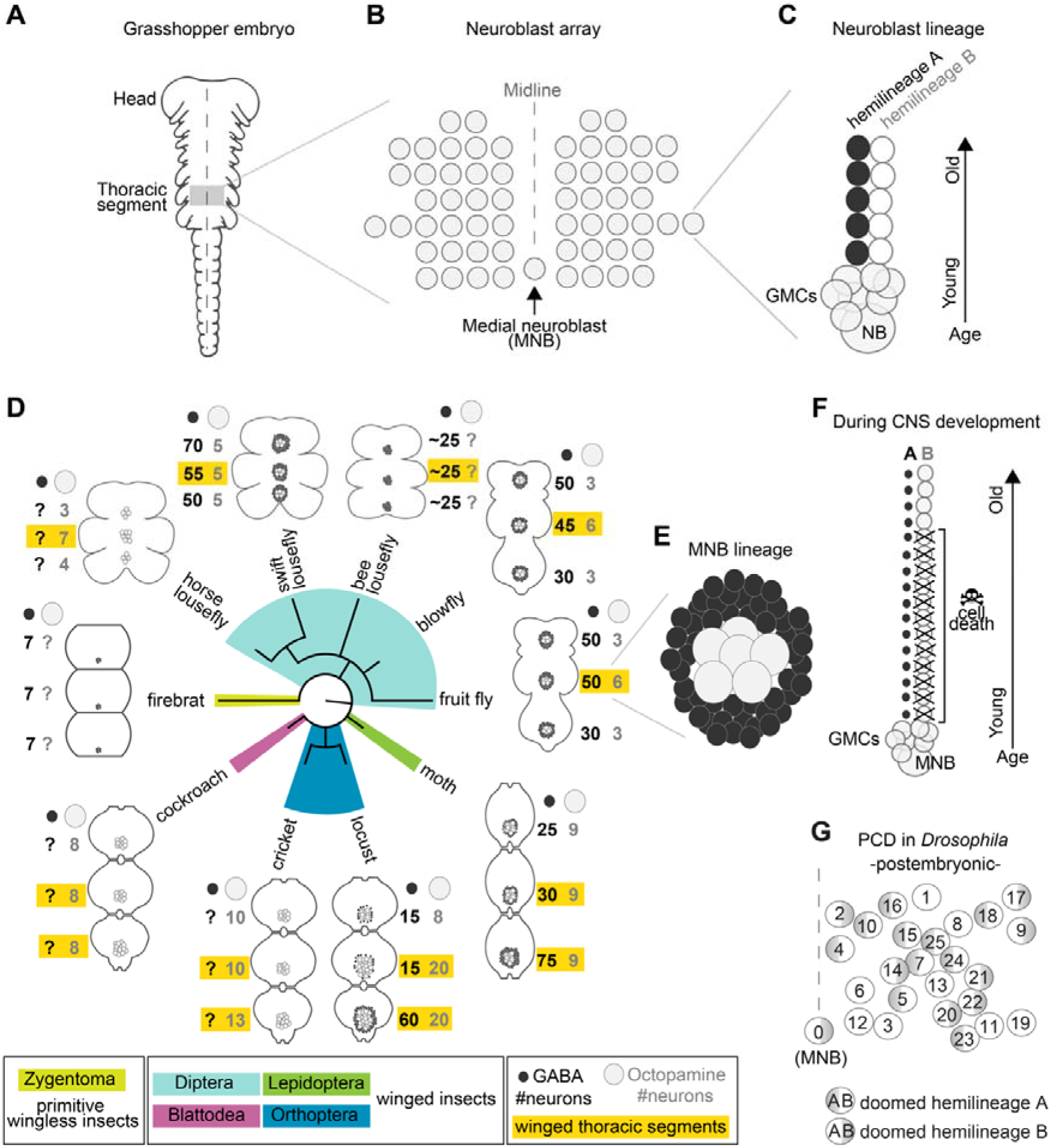
Median neuron numbers vary between insect species. **(A)** Cartoon of a young grasshopper embryo, here used as a depiction of a generic insect embryo (modified from Truman, 1996). Thoracic territory where one segments-worth of neuroectoderm generates an array of neuroblasts (grey box). **(B)** Schematic of neuroblast array showing bilaterally symmetric neuroblasts organised in 7 rows and 6 columns and a single median neuroblast (MNB) (modified from Truman, 1996). **(C)** Schematic of a lineage derived from a neuroblast (NB). Every NB buds off a ganglion mother cell (GMC) which undergoes a terminal division to generate 2 neurons with distinct cell fates, an A cell (black) and a B cell (grey). As the A and B cells result from a single division, one cannot be produced without the other. After several rounds of GMC divisions, a lineage produced by a single NB is composed of two half-lineages: ‘hemilineage A’ made up of all the A cells and ‘hemilineage B’ made up of B cells. Arrow indicates relative age, with newly-born cells located close to the NB. **(D)** Schematic showing summary of thoracic midline neuron population data from 5 orders of insects (colour coded in the phylogenetic tree and in the figure key bellow) including the primitive wingless firebrat *Thermobia domestica*, the cockroach *Periplaneta americana*, the cricket *Gryllus bimaculatus*, the locust *Schistocerca gregaria*, the moth *Manduca sexta*, the fruit fly *Drosophila melanogaster*, the blowfly *Calliphora vicina*, the horse lousefly *Hippobosca equina* and the flightless bee louse fly *Braula coeca* and swift lousefly *Crataerina pallida*. The numbers of GABAergic neurons (black cells) and octopaminergic neurons (grey cells) produced by the MNB is given for each thoracic segment. Except for the moth, a higher number of octopaminergic neurons can be found in winged segments in flying insects (yellow boxes). Cell numbers in this homologous lineage vary both between segments and species. Data on firebrats, cockroaches, crickets, locusts, moths, fruit flies and blowflies are compiled from Monastirioti et al., 1995; Stevenson and Spörhase-Eichmann, 1995; Witten and Truman, 1998; Schlurmann and Hausen, 2003; Lacin et al., 2019; and unpublished data from Dacks, Pflüger and Hildebrand (AM Dacks, personal communication, May 2020), while data on horse, swift and bee louseflies are from our own work. **(E)** Cartoon of *Drosophila* mesothoracic midline lineage populations with ‘hemilineage A’ cells revealed by GABA immunoreactivity (black) and ‘hemilineage B’ revealed by octopamine immunoreactivity (grey). **(F)** Schematic of MNB during development with ‘hemilineage A’ cells (black) and ‘hemilineage B’ (grey). The first neurons in hemilineage B cells survive (they are the oldest). After this point, all hemilineage B cells are removed by PCD whereas ‘hemilineage A’ cells from the same GMC division are left intact. **(G)** Schematic representation of the pattern of hemilineage-specific cell death in one hemisegment in the mesothorax. Each circle represents one lineage produced by one NB. Numbers represent postembryonic lineage nomenclature established by Truman et al. (2004). Shade circle refers to doomed hemilineage. Lineage 0 is the postembryonic name given to the MNB lineage.

In the *Drosophila* embryo a first wave of neurogenesis generates the larval nervous system (Booker and Truman, 1987; Truman and Bate, 1988) after which the majority of NBs become quiescent. Following reactivation from quiescence NBs produce neurons throughout larval life until the early pupal stages. These postembryonic neurons - which make up most of the adult CNS – extend simple neuritic processes into the neuropil and stall until the pupal-adult transition when they grow complex arborisations, synapsing with their target cells (Truman, 1990). In the VNC, NBs bud off a ganglion mother cell (GMC) which undergoes a terminal division to generate 2 neurons with distinctly different cell fates (an A cell and a B cell). As the A and B cells result from a single division, one cannot be produced without the other. After several rounds of GMC divisions, a lineage produced by a single NB is composed of two half-lineages: ‘hemilineage A’ made up of all the A cells and ‘hemilineage B’ made up of B cells (**Figure 1C**). Hemilineages act as functional units in adult flies (Truman et al., 2004, 2010; Lin et al., 2010; Harris et al., 2015; Shepherd et al., 2016, 2019; Lacin et al., 2019). For example, in the MNB lineage, hemilineage A cells mature into GABAergic local interneurons while hemilineage B cells become efferent octopaminergic neurons (**Figure 1F**). Our previous work showed that a common fate of postembryonic neurons is PCD affecting approximately 40% of VNC hemilineages (**Figure 1G**) (Truman et al., 2010), this is also seen in the brain (Kumar et al., 2009; Bertet et al., 2014). The pattern of PCD is stereotypical and targets the same hemilineages across individuals. Taken together, the breadth of PCD suggests its major role in shaping the final makeup of the adult nervous system, while its stereotypy points towards a heritable genetic basis. We therefore propose that changes in neural circuits may results from heritable alterations in the extent and pattern of PCD in hemilineages.

Here we show an extensive role for PCD in patterning the insect CNS, from primitive firebrats to the most derived true flies. We see evidence of hemilineage-specific PCD in primitively wingless firebrats and hippoboscid louseflies, suggesting that it is deployed widely. We search for changes in hemilineage programs of cell death during the evolution of flightlessness in true flies *Braula coeca* (the bee louse) and *Crataerina pallida* (the swift louse), and show that PCD is likely responsible for reductions in flight hemilineages within thoracic networks. Mimicking an evolutionary role for reducing PCD, we show that blocking PCD in the octopaminergic hemilineage produced by the MNB in *Drosophila melanogaster* results in mature differentiated ‘undead’ neurons that survive into adulthood, elaborate complex arborizations and are functional. Using new tools, we also show that this PCD takes place early, very soon after neurons are born. Our work highlights the importance of viewing hemilineages as functional units of neurodevelopment in all insects and shows that their alteration through an early mode of PCD can lead to adaptive changes in central circuits during evolution.

## RESULTS

### Early mode of neuronal cell death is found in many different insects

Because PCD selectively eliminates hemilineages in the fruit fly, and has been reported to kill off immature octopaminergic neurons produced by the MNB in the grasshopper (Jia and Siegler, 2002), we wondered if PCD occurs in the development of the CNS in a ‘primitive’ wingless insect. Using TUNEL labelling in the firebrat *Thermobia domestica* (**Figure 2A,B**), we found dying cells close to many NBs in all thoracic neuromeres at 50-55% of embryonic development (**Figure 2C,D,E**). We also looked for evidence of PCD during VNC development in the flightless dipteran *Crataerina pallida*, the swift lousefly (**Figure 2F**)*. Crataerina* is a viviparous haematophagous ectoparasite of the swift *Apus apus* (Hagan, 1951; Bequaert, 1952; Hutson, 1984; Walker and Rotherham, 2010a; Walker and Rotherham, 2010b). We focused on the swift lousefly, firstly to ascertain if the loss of flight during evolution resulted in increased PCD of flight hemilineages, and secondly, as it is a dipteran fly, we anticipated that we could use our specialist knowledge and immunohistochemical tools established in *Drosophila*.

**Figure 2.**
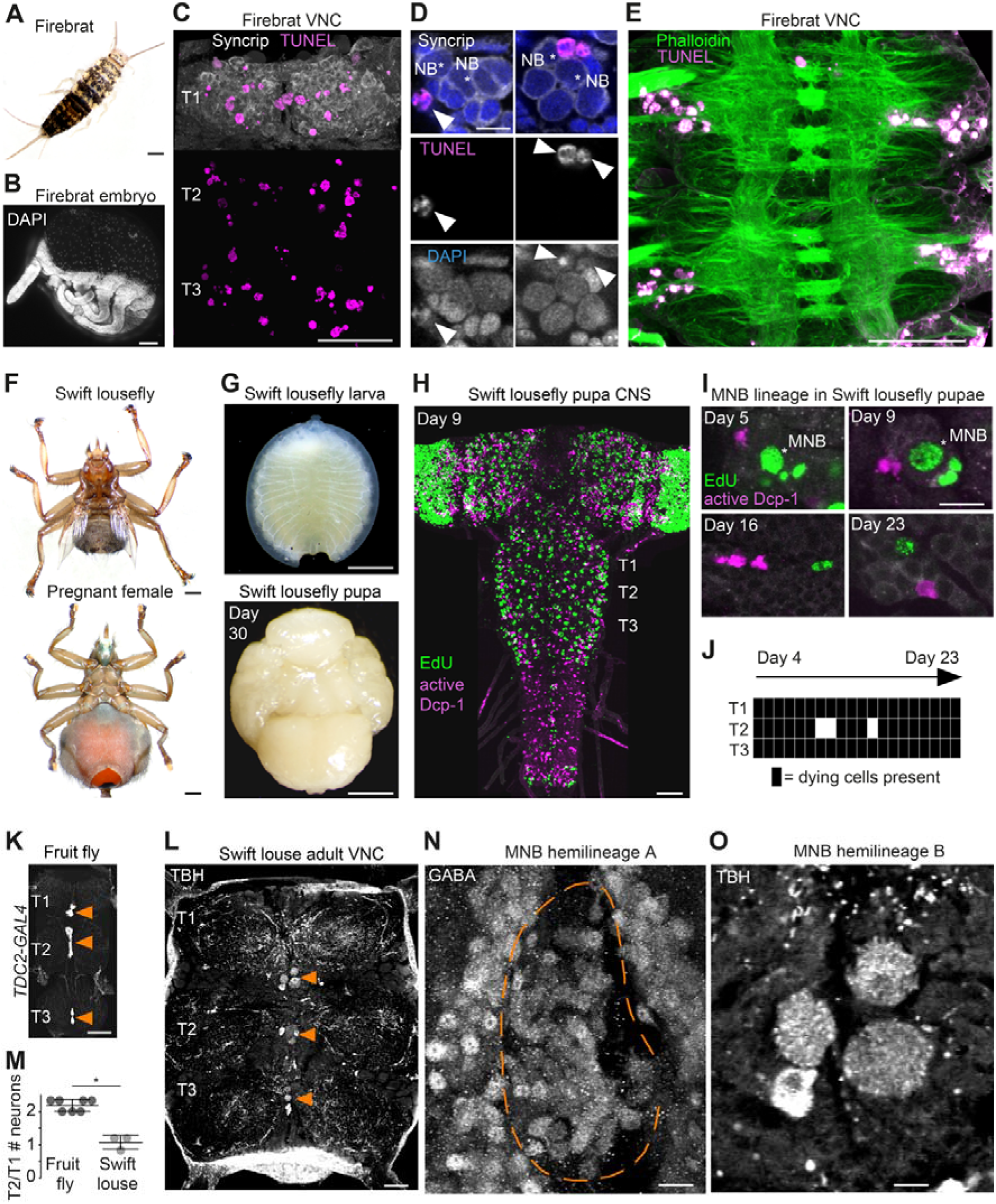
Cell death during neurogenesis is found in many insects. **(A)** Adult firebrat. Scale bar, 1mm. **(B)** Maximum intensity projection of DAPI staining in a wholemount firebrat embryo (*Thermobia domestica*) at 50-55% of embryonic development. Scale bar, 100 μm. **(C)** Dying cells in the thoracic VNC of a firebrat embryo labelled using TUNEL (magenta) and Syncrip (white) antibodies. Syncrip was used here as a proxy for Neuroglian staining to reveal lineages. Scale bar, 50 μm. **(D)** Dying cells (arrowheads) are located close to NBs (*). Scale bar, 10 μm. **(E)** Dying cells (magenta, TUNEL) are located in the cortex of the VNC, where neurogenesis takes place, and not in the neuropil (green, Phalloidin) in a firebrat embryo. Scale bar, 50 μm. **(F)** Dorsal view of an adult swift lousefly with vestigial wings (top). Ventral view of a female pregnant with a prepupa (bottom). Scale bars, 1 mm. **(G)** Swift lousefly larva (top) and day 30 swift lousefly pupa removed from its puparial case (bottom). Scale bars, 1 mm **(H)** EdU labels proliferating cells and antibody-labelling for active Dcp-1 reveals dying cells in the CNS of a swift lousefly pupa 9 days after pupariation. Scale bar, 50 μm. **(I)** Dying cells in lineage 0 labelled with antibodies for active Dcp-1 are located close to proliferating cells (e.g., NB*) throughout neurogenesis at Day 5 (top left), Day 9 (top right), Day 16 (bottom left) and Day 23 (bottom right) after pupariation. Scale bar, 10 μm. *n* = 1 each. **(J)** The occurrence of active Dcp-1-positive cells in lineage 0 in T1, T2 and T3 from Day 4 to Day 23 after pupariation in swift lousefly pupae (*n* = 1 each). Each black box indicates one occurrence. **(K)** Wild-type octopaminergic neurons in hemilineage 0B in a *Drosophila melanogaster* VNC labelled with CD8::GFP driven by *TDC2-GAL4* (orange arrowheads). Scale bar, 50 μm. **(L)** Octopaminergic neurons in hemilineage 0B in a swift lousefly VNC labelled with antibodies for tyramine β-hydroxylase (TBH, orange arrowheads). Fluorescence in the neuropil is derived from secondary antibodies trapped in the tracheal system and does not mark the true presence of TBH protein. Scale bar, 50 μm. **(M)** Quantification of T2/T1 number of octopaminergic neurons in fruit flies and swift louseflies shows that swift louseflies have lost the T2-specific higher numbers typical of flying dipterans (*P*\* = 0.012, Mann-Whitney. *n* = 7 fruit flies, *n* = 3 swift louseflies). Bars represent mean ± standard deviation. **(N)** Cluster of cell bodies belonging to hemilineage 0A (dashed outline) labelled with antibodies for GABA and **(O)** cell bodies belonging to hemilineage 0B labelled with TBH antibodies in the prothorax (T1) of a swift lousefly.

The swift lousefly is viviparous, females carry a single larva inside their abdomen where it feeds on ‘milk’ from fatty glands until pupariation, when the mother deposits the prepupa just before hardening of its pupal case (**Figure 2F,G**). Similar to the tsetse fly (Truman, 1990), neurodevelopment is significantly delayed compared to typical dipteran flies-the nervous system only acquires dipteran larval features many days after pupariation (**Figure 2H**). Using EdU to label proliferating cells and immunostaining for active Dcp-1 (**Figure 2H,I,J**), we found dying cells located close to NBs throughout the 24 days of pupal neurogenesis (manuscript in preparation). In the MNB lineage, which is easily identified by its medial position and projection pattern in the neuropil, we found cell death in thoracic segments at all time points examined, from day 4 after pupariation to day 23 (**Figure 2I,J**).

Similar to cockroaches, which are not efficient fliers (Eckert et al., 1992), the flightless swift louseflies have the same number of octopaminergic neurons in each thoracic segment (see **Figure 1D**). In flying insects these octopaminergic neurons are required for flight maintenance and initiation (Roeder, 2005) and display variability, with more neurons found in winged segments (Stevenson and Spörhase-Eichmann, 1995). We labelled lousefly octopaminergic neurons from hemilineage 0B using antibodies for tyramine β-hydroxylase (Monastirioti et al., 1996) and by comparing the ratio of octopaminergic cells in the mesothorax (winged segment) and the prothorax (lacks wings), we found that, unlike flying dipterans, the swift lousefly has lost segment-specific variability of cell numbers (fruit fly [2.2±0.2, n = 7] versus swift lousefly [1.1±0.2, n = 3], p = 0.012, Mann-Whitney U = 0, Mann-Whitney t-test) (**Figure 2K,L,M,N,O**). In the sister hemilineage 0A we found a considerably larger number of GABAergic neurons (**Figure 2N**), suggesting that PCD is responsible for the selective elimination of the octopaminergic hemilineage.

Our observations in firebrats and louseflies suggest that PCD during neurogenesis may be a universal and ancestral feature that sculpts the nervous system of all insects. To further explore if changes in PCD have been deployed during evolution to accommodate adaptive modifications to behaviour, we next searched for evidence of increased PCD in other flight hemilineages of flightless dipterans.

### Changes in hemilineage programs of PCD in flightless dipterans suggest adaptive modifications to neural circuits

To explore the possibility that changes in the pattern and/or extent of PCD are adaptive, we looked for evidence of evolutionary modifications in the VNC of other species. We specifically asked whether lineages known to function in flight circuitry might be modified in flightless insects. Using antibodies for Neuroglian, we compared homologous hemilineages involved in the flight circuits of two flightless dipterans, the swift lousefly *Crataerina pallida* and the bee lousefly *Braula coeca* (**Figure 3A,B,C,D**). *Braula*, a close relative of drosophilids, is wingless, lacks halteres and has an extremely reduced thorax (**Figure 3A**). Bee louseflies spend their entire adult life as kleptoparasites on the honeybee *Apis mellifera* (Imms, 1942; McAlister, 2018).

**Figure 3.**
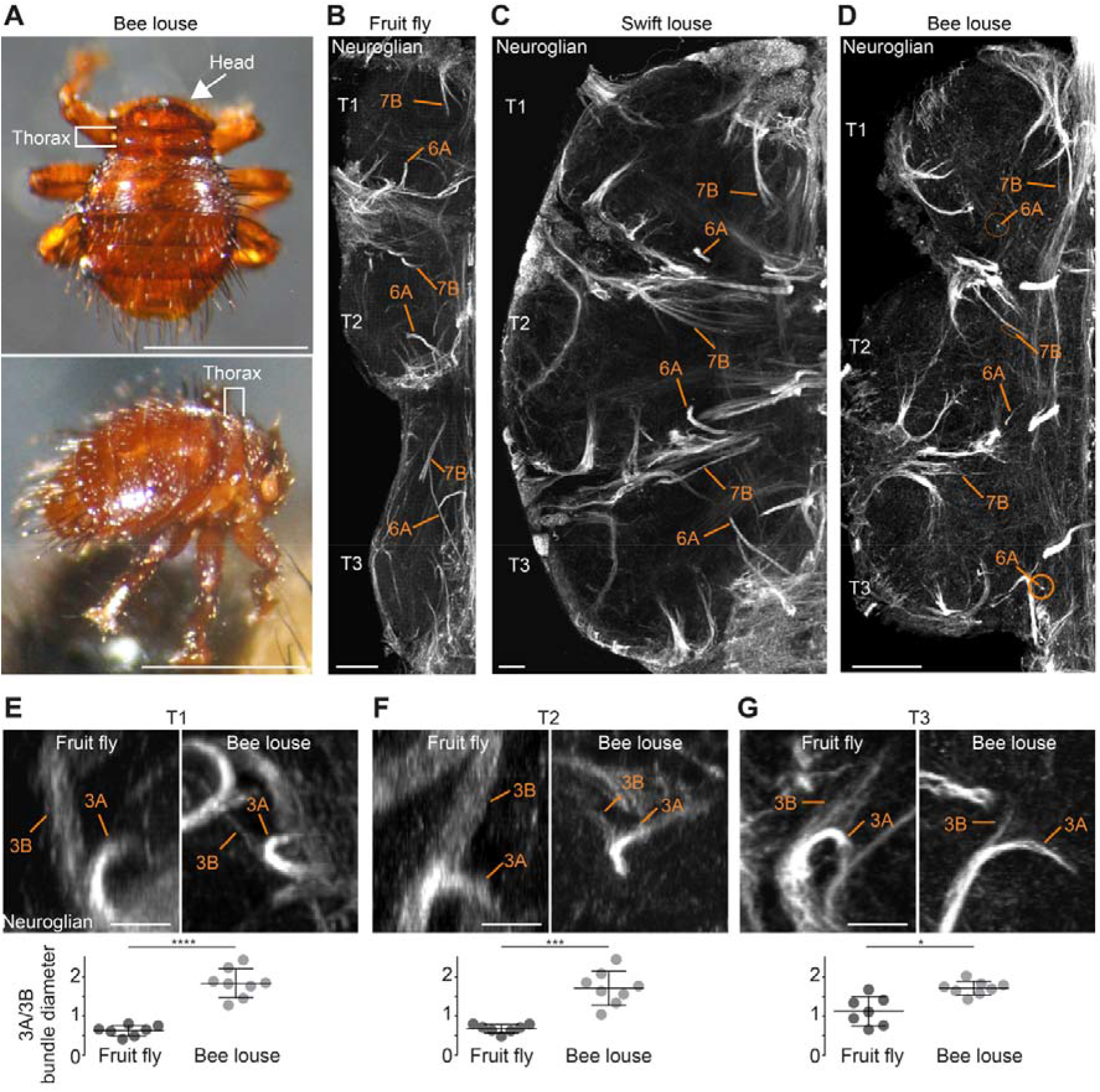
Neuronal cell death sculpts the wing circuitry of wingless dipterans. **(A)** Dorsal (top) and side view (bottom) of an adult bee lousefly with a reduced thorax lacking wings and halteres. Scale bar, 1 mm. (**B-D**) Hemilineage VNC fiber tracts labelled with Neuroglian in a fruit fly (**B)**, a swift lousefly (**C**) and a bee lousefly (**D**). Shown are hemilineages 6A and 7B which are reduced in the bee lousefly in all three thoracic segments (T1, T2, T3). Scale bars, 50 μm. (**E-G**) 3A and 3B hemilineage fiber tracts labelled with Neuroglian in a fruit fly (**top left**) and a bee lousefly (**top right**). Shown are maximum intensity projections, chosen to best display hemilineages, from cross-section T1 (**E**), T2 **(F)** and frontal perspectives T3 **(G)**. Quantifications of 3A/3B hemilineage bundle diameter ratios in fruit flies and bee louseflies are given below (*P*\*\*\*\* < 0.0001 in T1, independent samples t-test, *P*\*\*\* = 0.0001 for T2, Welch’s t-test, *P*\* = 0.0044, Welch’s t-test. *n* = 7 fruit flies each, *n* = 8 bee louseflies each). Bars represent mean ± standard deviation.

Comparing the neuroglian-labelled axon bundle width of homologous hemilineages between flightless and flying species, we found that hemilineages 3B, 5B, 6A, 7B, 11B, 12A and 19B, which innervate the wing neuropil (Shepherd et al., 2019), are reduced in bee louseflies, but not in swift louseflies (**Figure 3B,C,D**). Among these, 3B, 6A, 11B, 12A and 19B belong to lineages in which both hemilineages survive in fruit flies (see schematic in **Figure 1G**). A difference in axon bundle diameter between sister hemilineages could indicate a difference in cell number, possibly established by PCD during development. We chose to quantify the ratio of axon bundle diameters in lineage 3 because we expected a reduction in hemilineage 3B, which innervates the wing neuropil, but no change in hemilineage 3A, which projects into the leg neuropil (Shepherd et al., 2019). The fiber tracts of 3A and 3B originate as a common bundle and split only in the intermediate neuropil. After they split, the sister fiber tracts sit in the same plane before they defasciculate, making them easy to trace and compare (**Figure 3E,F,G**). We found that the ratio between sister hemilineages A and B is significantly higher in bee louseflies compared to fruit flies, indicating that hemilineage 3B, which controls flight-related behaviours, is severely reduced in these flightless flies (T1: fruit fly [0.6±0.1, n = 7] versus bee louse [1.8±0.4, n = 8], p < 0.0001, t = −8.084, independent samples t-test; T2: fruit fly [0.7±0.1, n = 7] versus bee louse [1.7±0.4, n = 8], p = 0.0001, F = 42.22, Welch’s t-test; T3: fruit fly [1.1±0.4, n = 7] versus bee louse [1.7±0.2, n = 8], p = 0.0044, F = 14.84, Welch’s t-test). Even though we cannot make a precise inference of cell numbers in each hemilineage, as the fiber tract of hemilineage 3B appears frayed while 3A is more compact in both species, our results clearly show there has been an adaptive reduction of 3B, associated with the loss of flight machinery.

### Blocking programmed cell death in *Drosophila* generates identifiable, differentiated populations of undead neurons

After observing the outcome of increased PCD in hemilineages consistent with adaptations to the loss of flight in bee louse and swift louse, we wondered if, conversely, by only reducing PCD, we could generate novel functional expansions of a hemilineage. To explore this, we made use of the powerful genetic tools available in *Drosophila* to block PCD in the MNB lineage to determine if ‘undead’ cells survive into adulthood, elaborate their neurites and acquire a distinctive neurotransmitter identity.

From our previous work (Truman et al., 2010), we know that during postembryonic neurogenesis MNB hemilineage A survives, expresses Engrailed (Truman et al., 2004) and differentiates into GABAergic interneurons (Lacin et al., 2019). Because during embryonic development the MNB hemilineage B produces a small number of octopaminergic neurons, we hypothesised that preventing PCD would generate additional octopaminergic neurons in the later postembryonic phase of neurogenesis. In postembryonic nomenclature, neurons generated by the MNB are collectively called lineage 0 and therefore we will refer to octopaminergic neurons generated by the MNB as hemilineage 0B.

Using the octopaminergic neuron driver, *TDC2-GAL4,* we observed a 4 to 9-fold increase in the number of octopaminergic neurons in the thoracic VNC of *H99/XR38* adult flies deficient for proapoptotic genes (*hid^+/−^*, *grim^+/−^*, *rpr^−/−^* and *skl^+/−^*) (White et al., 1994; Peterson et al., 2002) compared with wild-type control animals (**Figure 4A,B,C,D**), (T1: 20.9 ± 2.3, Mann-Whitney U = 0, p = 0.0002; T2: 26.3 ± 4.4, Mann-Whitney U = 0, p = 0.0004; T3: 27.5 ± 3.5, Mann-Whitney U = 0, p = 0.0004; n = 11 each). These ‘undead’ neurons also express the vesicular glutamate transporter VGlut (**Figure 4E,F**), just like wild-type octopaminergic neurons (Greer et al., 2005).

**Figure 4.**
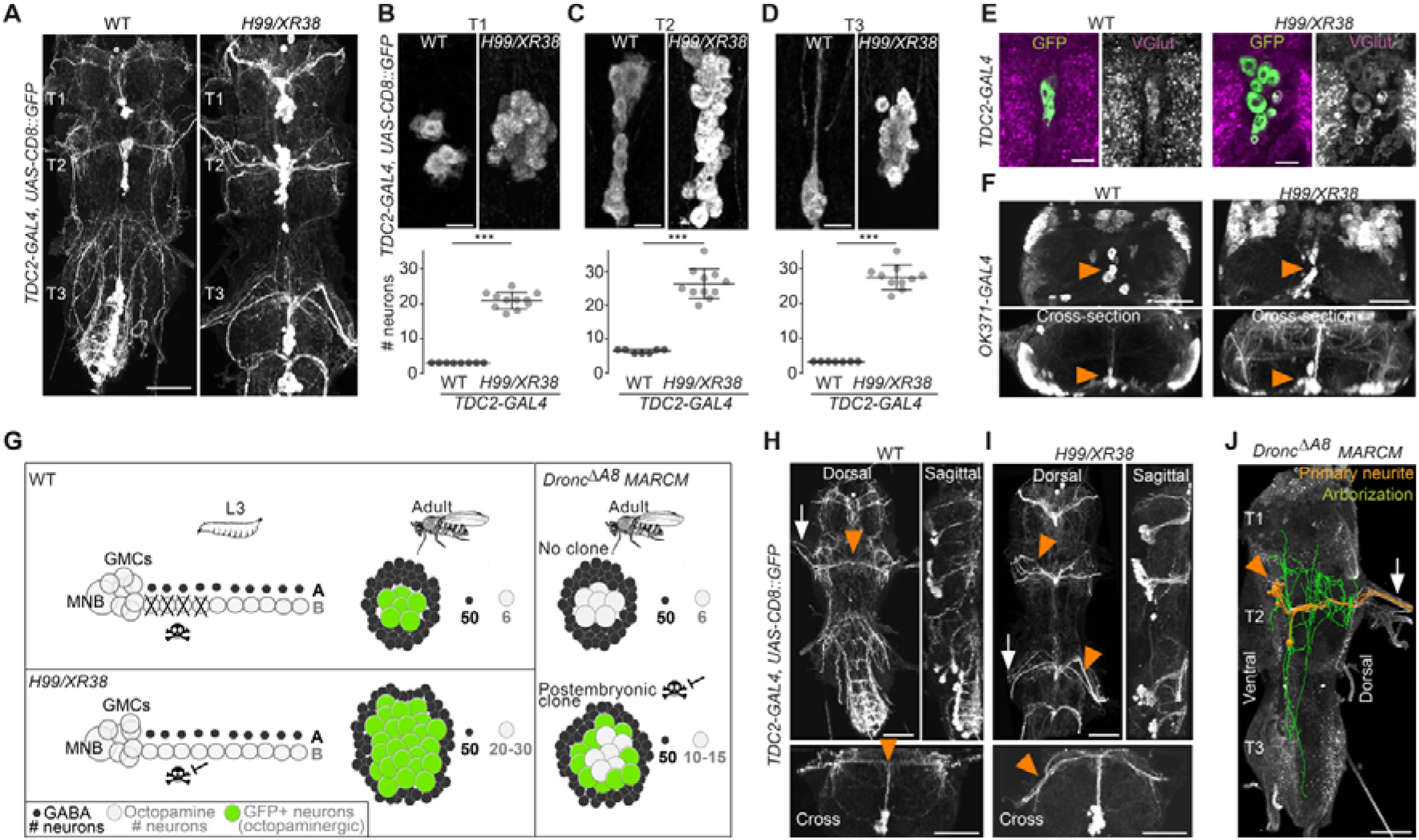
Blocking death results in differentiated undead neurons in the medial neuroblast lineage. **(A)** CD8::GFP expression driven by *TDC2-GAL4* in octopaminergic neurons from hemilineage 0B in the VNC of wild-type (WT, left) and PCD-blocked adult flies (*H99/XR38* deficient for *hid^+/−^*, *grim^+/−^*, *rpr^−/−^* and *skl^+/−^*, right). Scale bar, 50 μm. **(B-D)** Quantifications of the number of *TDC2-GAL4*-positive octopaminergic neurons in the VNC of WT and *H99/XR38* adult flies. Bars represent mean ± standard deviation. (**B**) \*\*\**P* = 0.0002 in T1, **(C)** \*\*\**P* = 0.0004 in T2, **(D)** \*\*\**P* = 0.0004 in T3, Mann-Whitney. Scale bar, 10 μm. *n* = 11 each. Mann-Whitney. Scale bar, 10 μm. *n* = 11 each. **(E)** Antibodies for the vesicular glutamate transporter VGlut and **(F)** GFP expression driven by the glutamatergic driver line *OK371-GAL4* label both WT and undead (*H99/XR38*) octopaminergic neurons. Orange arrowheads indicate cell bodies. Scale bars, 10 μm (VGlut), 50 μm (OK371). **(G)** Schematic of TDC2-GAL4 expression in postembryonic lineage 0 in wildtype and *H99/XR38* third instar larvae and adults (left panels). Postembryonic hemilineage B populations only start expressing TDC2-GAL4 in early pupal development and maintain it throughout adult life. MARCM mosaic clones that are homozygous for a null *Dronc* allele lack GAL80. These show robust expression of GAL4 in small numbers of surviving postembryonic hemilineage B cells (right panel). In adult WT and *H99/XR38* flies, GFP is expressed in both embryonically-born and postembryonic TDC2-positive neurons, while in MARCM flies GFP is *only* present in postembryonic cells. **(H-I)** CD8::GFP expression driven by *TDC2-GAL4* in WT (**G**) and *H99/XR38* (**H**). WT and undead primary neurites project dorsally and branch extensively in the dorsal neuropil. In WT neurons the primary neurite bifurcates at the dorsal midline, while undead neurons are unable to bifurcate and turn to one side (orange arrowheads). In *H99/XR38* flies which contain both WT and undead neurons, the primary neurite to one side is thicker (orange arrowhead). Both WT and undead neurons join thoracic nerves (white arrows). Scale bars, 50 μm. **(J)** Reconstructed arborizations of undead neurons expressing CD8::GFP driven by *TDC2-GAL4* in flies bearing MARCM clones homozygous for the loss of function allele *dronc*Δ*A8* (in which PCD is blocked). The 3D-rendered image is tilted at a 45° angle. Undead neurons have somata that are located at the ventral midline (orange arrowhead), branch extensively in the neuropil (green), have a turning primary neurite (orange), and project to the periphery through a thoracic nerve (arrow). Scale bar, 50 μm.

Ideally, to label and manipulate dying neurons from hemilineage 0B, we require a specific driver line expressed only in the newly-born doomed neurons. To test if we could use *TDC2-GAL4* to label and manipulate dying neurons from hemilineage 0B during their development, we performed a timeline of expression in wild-type and *H99/XR38* flies (**Figure S1**). Unfortunately, even though undead neurons are generated from L2 onwards in *H99/XR38* flies, the *TDC2-GAL4* is only active in the undead cells days later. Gradually, in pupae, TDC2-GAL4 expression reveals the remaining undead B cells (**Figure S1C,D**). We concluded that the TDC2 driver line cannot be used to visualize and manipulate newly-born postembryonic ‘doomed cells’. Instead, TDC2-GAL4 allowed us to accurately reveal ‘undead’ hemilineage 0B neurons *but* only in the adult (see cartoon **Figure 4G**).

To ensure sparse labelling and precise manipulation of doomed cells from hemilineage 0B, we generated postembryonic *TDC2-GAL4*-expressing MARCM clones homozygous for the loss-of-function allele *Dronc*^*ΔA8*^(in which PCD is inhibited) (Kondo et al., 2006; Truman et al., 2010). This strategy guarantees that, even though cell death can be rescued in other lineages, it is only *TDC2*-positive postembryonic neurons that express UAS-based tools.

Analysis of the projection patterns of undead neurons revealed that they display both common and distinct features compared to their wild-type embryonically-born counterparts. Similar to wild-type octopaminergic cells (Monastirioti et al., 1995), the undead neurons have cell bodies located ventrally at the midline, at the posterior border of the thoracic segment (**Figure 4H,I,J**), project a primary neurite in the dorsal-most region of the neuropil, the tectulum (Court et al., 2017), and join thoracic nerves (Pauls et al., 2018) (**Figures 4H,I,J** and **S2**). Unlike wild-type cells which bifurcate and branch extensively in the tectulum, the primary neurite of undead neurons fails to bifurcate, branches in both dorsal and ventral regions of the neuropil and sends projections to neighbouring segments (**Figures 4J** and **S2**). As we describe in **Figure S1**, a few wild-type octopaminergic neurons are produced in all thoracic segments during postembryonic neurogenesis in lineage 0. We propose that the very few bilateral projecting neurons we encounter in our clones are wild-type cells. To avoid any uncertainty when performing our behavioural experiments (below), we excluded flies which contained a bifurcating neuron in undead MARCM clones. Thus, using MARCM clonal approaches we show that undead neurons in hemilineage 0B become octopaminergic, elaborate complex neurites and join thoracic nerves.

### Undead neurons are functional and integrate into motor networks

We next asked if these differentiated undead neurons are functional. To address this, we tested if activating undead neurons with the warm temperature-gated ion channel TrpA1 in headless adult *Drosophila* could elicit behaviours (**Figure 5 and Video S1**). In *TDC2-GAL4* positive control flies (expressing TrpA1 in wild-type embryonic-born octopaminergic neurons), thermogenetic stimulation induced long bouts of locomotion (**Figure 5C,D** and **Video S1**). Importantly, we found *UAS-TrpA1* negative controls (in the absence of a GAL4) and MARCM control flies (containing no GAL4-positive clones) did not walk in response to temperature elevation (**Figure 5B,C,D** and **Video S1**). We found that the activation of undead neurons expressing TrpA1 caused decapitated males to walk (**Figures 5B,C,D,E** and **Video S1**). These data are consistent with the observation that octopamine applied to the exposed anterior notum of decapitated flies causes walking (Yellman et al., 1997) and suggests that undead neurons are functional and capable of releasing neurotransmitters in the CNS. The extent of walking was greater in positive control flies than in the undead MARCM condition (compare panels in **Figure 5C** and **Video S1**). This is likely because, alongside activating octopaminergic neurons in the VNC, in positive controls we also stimulated the severed axons of octopaminergic cells in the brain which send descending projections to the VNC.

**Figure 5.**
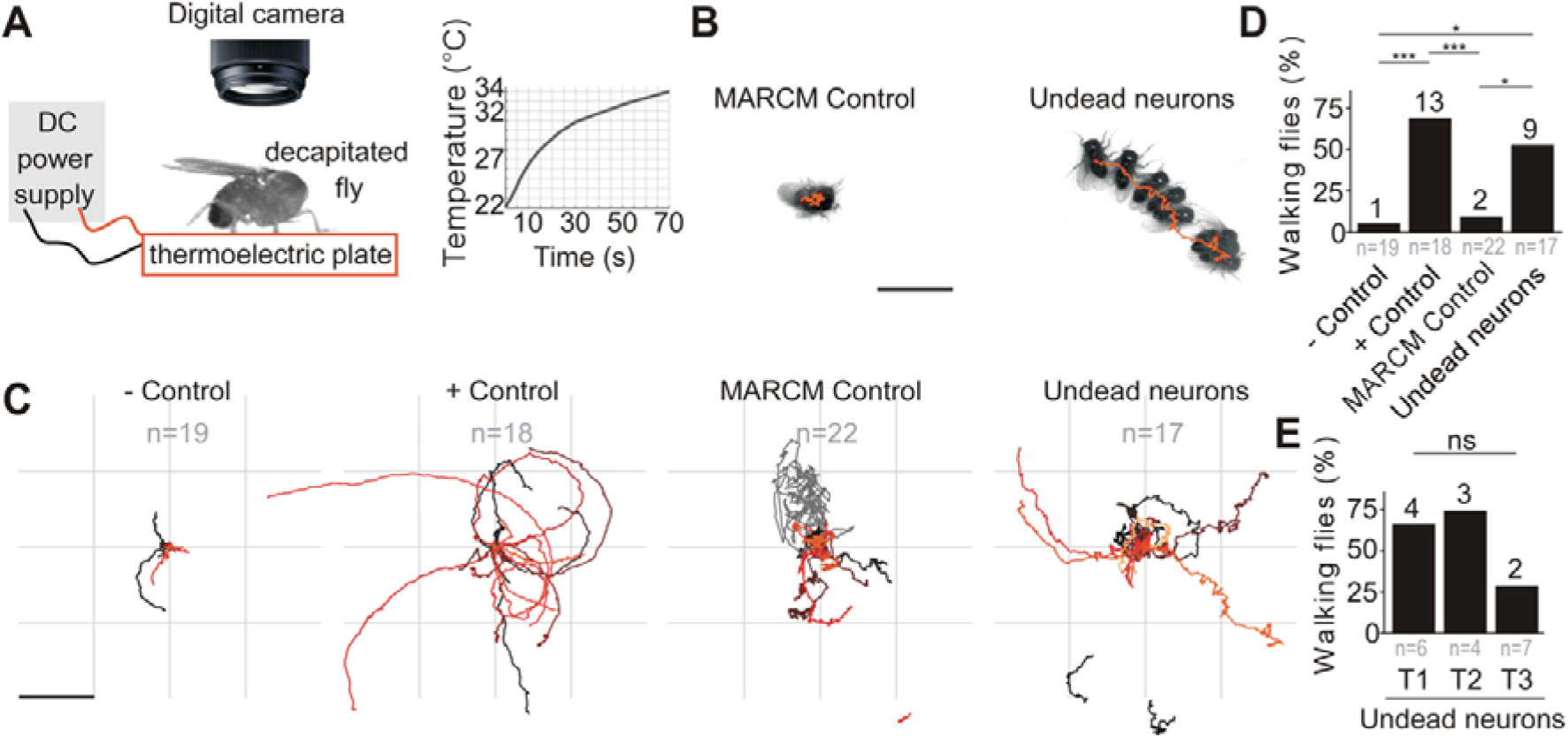
Undead neurons are functional: thermogenetic activation of undead neurons induces walking in decapitated *Drosophila*. **(A)** Schematic of behavioural assay with TrpA1 activation. Decapitated flies are placed on a thermoelectric plate connected to a DC power supply, exposed to a temperature ramp (right panel), and filmed from above using a digital camera. **(B)** Examples of a stationary MARCM Control fly and a walking fly with undead neurons. Images represent maximum intensity projections of 13 frames at 0.3 fps tracing the centroid over time (orange line). Scale bar, 5 mm. **(C**) Fly body tracks generated by identifying the geometric centre of the fly body in each frame and storing the centre coordinates, plotted as a continuous line, one for each fly (walking or stationary) for negative controls (*UAS-TrpA1*), positive controls (*TDC2>TrpA1*), MARCM control flies and flies with MARCM clones of Undead neurons. Each trace represents one individual fly. Scale bar, 5 mm. **(D)** Quantification of the percentage of flies that walked per experimental group. *P*\*\*\* = 0.0002 for negative versus positive control, *P*\*\*\* = 0.0002 for positive controls versus MARCM Control, P* = 0.0242 for MARCM control versus undead neurons, *P*\* = 0.0135 for negative control versus undead neurons, Pearson’s chi-squared corrected for multiple comparisons using a Bonferroni correction. *n* = 19 for negative controls, *n* = 18 for positive controls, *n* = 22 for MARCM control, *n* = 17 for undead neurons. *n* numbers for each group are given below and the number of flies which walked is shown above each bar. **(E)** Quantification of the number of walking undead neuron flies split into 3 anatomical subgroups according to the location of MARCM clones in T1, T2, or T3. *P*^ns^ = 0.2628, Pearson’s chi-squared. *n* = 6 for T1, *n* = 4 for T2, *n* = 7 for T3. Numbers at the base are the number of walking flies. The percentage is shown above each bar.

To determine if undead neurons are integrated into thoracic motor circuits, we recorded the activity of mixed undead and wild-type octopaminergic neuron populations expressing GCaMP6s (an activity reporter) and tdTomato (an anatomical fiduciary) in intact *H99/XR38* flies during tethered behaviour on a spherical treadmill (Chen et al., 2018) (**Figure 6A,B**). In these animals we observed conspicuous increases in neural activity during air-puff-induced walking in both wild-type controls (**Figure S3**) and in *H99/XR38* flies (**Figure 6C,D,E** and **Videos S2-4**). Because undead neurons outnumber their wild-type counterparts by a ratio of 6.5 to 1, these results imply that both neuronal types are active in *H99/XR38* flies. Supporting this, we observed an increase in GCaMP6s fluorescence across all subregions along the width of the primary neurite bundle (**Figure S4** and **Video S4**). Taken together these data reveal that ‘undead’ neurons in the adult fly are functional and integrate into motor networks.

**Figure 6.**
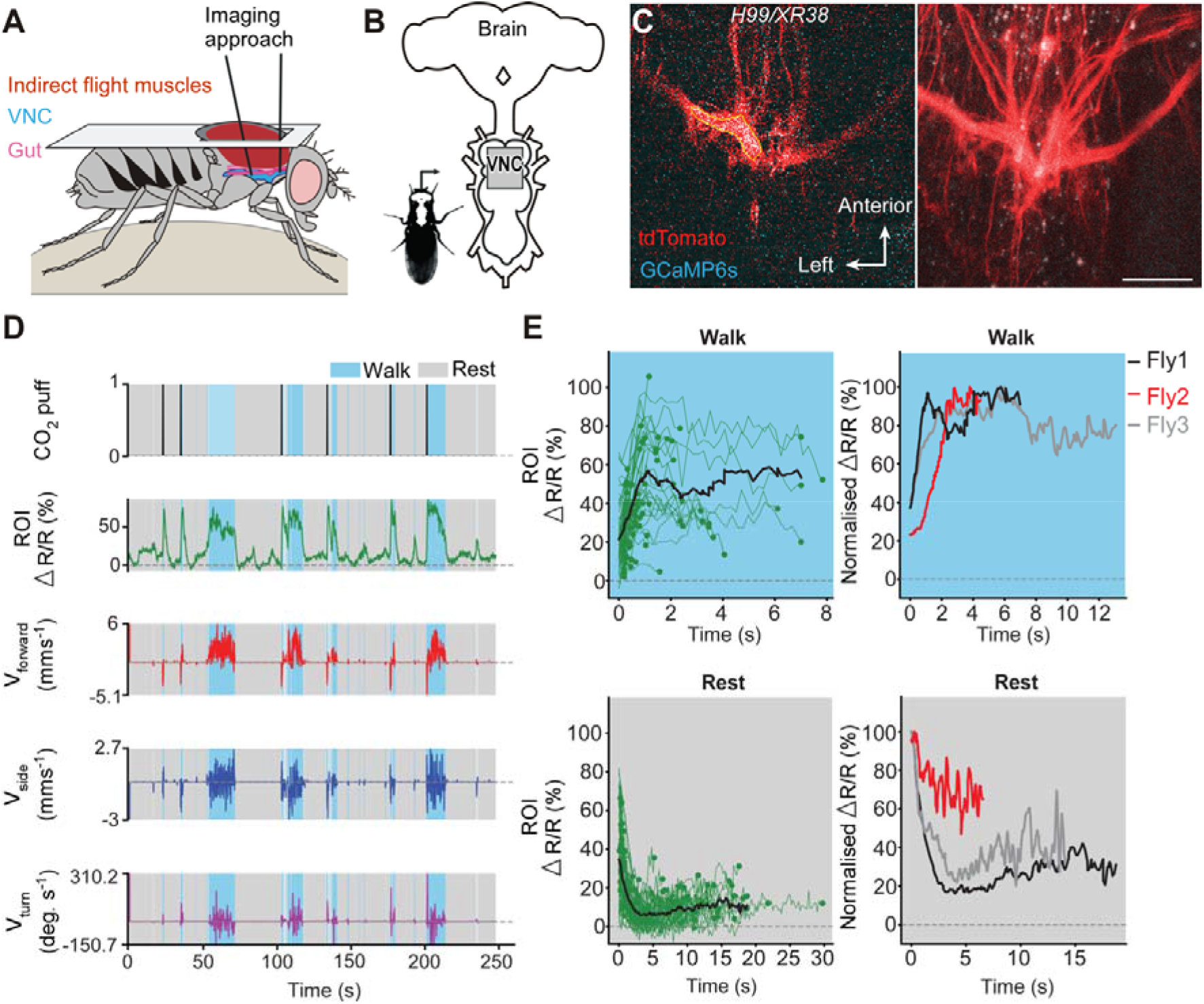
Undead neurons integrate into VNC networks: undead neurons are active during naturalistic behaviours in intact adult *Drosophila*. **(A)** Schematic of the dorsal thoracic dissection and approach for ventral nerve cord functional imaging in tethered, adult flies. **(B)** Location of the imaging region of interest (grey box) with respect to a schematic of the adult CNS. **(C)** Raw 2-photon image of *TDC2-GAL4-positive* neurons co-expressing tdTomato (red) and GCaMP6s (cyan) in *H99/XR38* flies (**left**). Region-of-interest used to calculate %ΔR/R is outlined (yellow). Standard deviation z-projection of a dorsal-ventral image stack of the functional imaging region-of-interest in **b** (**right**). Scale bar, 50 μm. **(D)** Representative behavioural and functional imaging data in *H99/XR38* flies. Shown are: CO_2_ stimulation (black), %ΔR/R (ratio of GCaMP6s / tdTomato) signal (green), and ball rotations indicating forward walking (red), sideways walking (blue), and turning (purple). The behaviour of the fly was classified as either walking (light blue), or resting (gray) by applying a threshold on ball rotation speed. **(E) (left)** Individual (green) and average (black) %ΔR/R traces within each behavioural epoch for walking (n = 82) and resting (n = 86) events processed from 750 s of imaging data. Solid green circles indicate the end of a behavioural epoch. The average trace (black line) was calculated for only periods with 4 or more traces. (**right**) Normalised average %ΔR/R traces for three different flies during walking and resting. The average (black) trace is the same as in the left panel.

### An early and rapid mode of developmental cell death eliminates significant numbers of newly-born neurons throughout postembryonic development in *Drosophila*

After observing the potential for blocking PCD to incorporate new neurons, we wondered which specific type of PCD is responsible, reasoning that our knowledge of how nervous systems evolve can only be enriched by pinpointing the exact developmental processes involved. The majority of studies on neuronal PCD in insects have focused on its role at metamorphic transitions, where it eliminates fully differentiated neurons either at puparium formation (Truman et al., 1994) or in adults post-eclosion (Kimura and Truman, 1990; Draizen et al., 1999). Both of these remodelling events are gated by ecdysteroids. However, finding dying cells close to NBs in firebrats and swift louseflies, combined with similar observations in *Drosophila* (Truman et al., 2010), made us consider that hemilineage-specific PCD takes place early, in newly-born neurons. This has been difficult to evaluate on a cell-by-cell basis within a complex nervous system.

To interrogate the postembryonic neuronal death we see here, we built a novel genetically encoded effector caspase probe called SR4VH (Figure 7A,B). SR4VH consists of a membrane-bound red fluorescent protein (Src::RFP) and a yellow fluorescent protein with a strong nuclear localisation signal from histone H2B (Venus::H2B) linked together by four tandem repeats of the amino acid sequence DEVD. When effector caspases cleave the DEVD site, Venus accumulates in the nucleus while RFP remains bound to the cell membrane (Figure 5B). This reporter is similar in design to Apoliner (Bardet et al., 2008), but has different subcellular localization signals as well as four tandem caspase cleavage sites instead of one. We also found that tethering the probe to the membrane with the myristoylation signal from Src means that there is no excess signal accumulation in the Golgi apparatus (Mukherjee et al., in prep). The nuclear localisation signal from H2B allows for highly efficient sequestration of cleaved Venus in the nucleus even in late stages of apoptosis, when the nuclear membrane is likely compromised.

**Figure 7.**
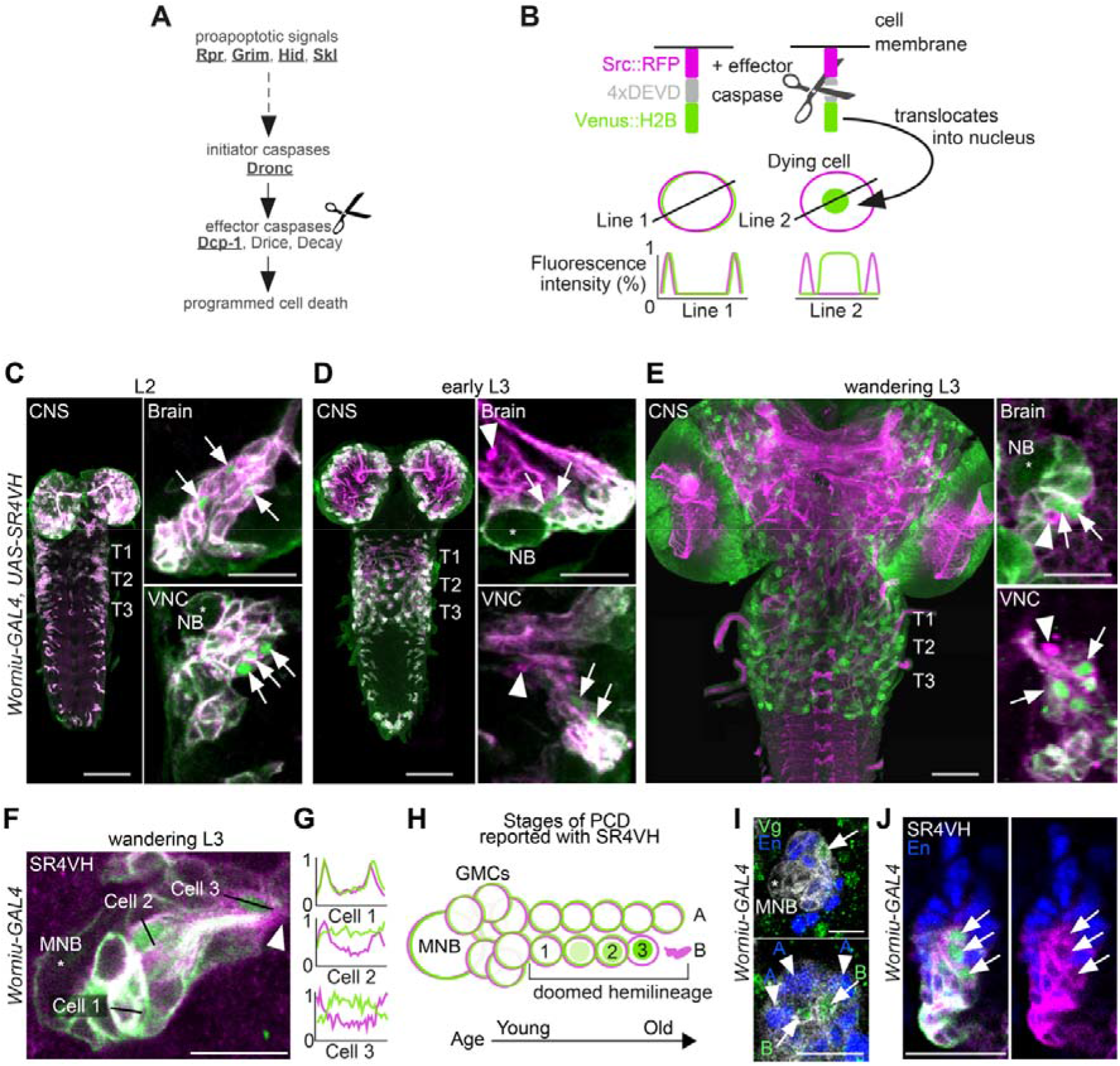
Early neuronal cell death occurs throughout postembryonic development. **(A)** Simple schematic of PCD in *Drosophila*. Elements disrupted in this study, or used as a PCD readout are underlined. **(B)** Schematic of effector caspase reporter SR4VH (top) and idealised fluorescence patterns in two cells with distinct caspase activity (bottom): RFP (magenta) and Venus (green) present in the cell membrane (Line 1) versus RFP at the cell membrane and Venus accumulation in the nucleus of a cell undergoing programmed cell death (PCD) (Line 2). **(C-E)** SR4VH driven by *Worniu-GAL4* reveals dying cells in the central nervous system (CNS) throughout postembryonic neurogenesis in a 2^nd^ instar (**C**, n = 9), early 3^rd^ instar (**D**, n = 8) and wandering 3^rd^ instar (**E**, n = 18) larva. Right panels show examples of lineages with dying cells located close to the NB (*) in the brain (top) and ventral nerve cord (VNC, bottom). Arrows indicate dying cell, arrowheads indicate RFP-positive dead cell membranes. Scale bars, 50 μm (left panels), 10 μm (right panels). **(F)** SR4VH driven by *Worniu-GAL4* reveals younger cells at earlier stages of cell death (Cell 2) are located closer to the MNB than older cells at later stages of cell death (Cell 3), which are closer to the lineage bundle (arrowhead). Image represents a single optical section. Scale bar, 10 μm. **(G)** Fluorescence intensity profiles (normalised to the maximum value along the line for each channel) plotted along the lines indicated in (**F)**. **(H)** Schematic representing successive stages of cell death correlated with distance from NB and cell age in doomed lineages as reported with SR4VH. **(I)** Revealing markers for lineage 0. Immature hemilineage A progeny revealed with anti-Engrailed (En, blue – arrowheads) and immature hemilineage B progeny labelled with anti-Vestigial (Vg, green – arrows) in a third instar *Worniu-GAL4; UAS-CD8::GFP* larva (white). Expression of markers is mutually exclusive. Scale bars, 10 μm. **(J)** SR4VH driven by *Worniu-GAL4* together with antibodies for Engrailed (En, blue) reveal that only Engrailed-negative cells from hemilineage 0B undergo PCD (white arrowheads) during postembryonic development in the thoracic VNC. Scale bar, 10 μm. *n* = 6.

Using the GAL4/UAS system and the neuroblast driver *Worniu-GAL4*, we found we could visualize postembryonic neurogenesis and label up to 20 of the most recently born progeny from a single NB (this is due to GAL4 and reporter perdurance). The number of progeny we can detect at any one time using *Worniu-GAL4* varies from 10 to 20, most likely as a result of differential proliferation rates across lineages. We confirmed that SR4VH is reliable as a reporter for cell death in larvae by analysing its expression pattern in all lineages of postembryonic neurons in the thoracic VNC and comparing it to our previous work on MARCM homozygous mutant clones of the initiator caspase *Dronc* (Truman et al., 2010). (**Figures 7C,D,E,F** **and** **S5**). We found dying cells associated with lineages in the brain and VNC throughout the whole of postembryonic neurogenesis (**Figures 5D,E,F**), which lasts for 3.5 days, from mid-2^nd^ instar (L2) to 12h after pupariation. As previously suggested (Truman et al., 2010), the time course of PCD indicates that cells die early – very soon after they are born – often before they have even extended a neuritic process. This death appears to be unlike the ‘trophic’ PCD found in vertebrates, where a neuron extends a process, interacts with its target cell and dies in the absence of appropriate survival signals. In support of an early onset of PCD, we were able to see sequential stages of cell death, dependent on the distance from the NB (**Figures 7G,H,I** **and** **S5B**). Older cells located further away from the NB appear to be at a more advanced stage of PCD indicated by the complete translocation of Venus from the membrane to the nucleus (Cell 3 in **Figure 7G,H**) and by the accumulation of RFP-positive dead cell membranes close to the lineage bundle (arrowheads in **Figure 7E,F**).

The number of dying cells within a doomed lineage varied from 1 to 8, with most lineages containing 1-2 dying cells from a total of 10-20 cells labelled with *Worniu-GAL4* (dying cells/lineage: 1.3 ± 1.5 given as average ± standard deviation; n = 444 doomed lineages from 5 VNCs). From a total of 444 doomed lineages, 243 harboured more than one dying cell, of which 148 displayed a progression of cell death (**Figure 7G,H**). Truman and Bate approximated the cell cycle of an NB to 55 minutes and that of a GMC to 6.5 hours, with 7 GMCs present in a proliferating lineage at all times (Truman and Bate, 1988). Therefore, after subtracting the NB and GMCs from clusters of 10-20 *Worniu-GAL4*-labelled cells, 2-12 will be neurons which resulted from 1-6 divisions, each separated in time by 55 minutes. This means that PCD was initiated early, at some time between 0 and 5.5 hours after neurons were born.

To look at death during the development of the MNB lineage (lineage 0) we imaged SR4VH in wandering L3 larvae and used molecular markers to identify members of hemilineage 0A and 0B. The transcription factors Engrailed/Invected (En/Inv) are known to be expressed in immature and fully differentiated interneurons of hemilineage A (Truman et al., 2004; Lacin et al., 2014, 2019; Allen et al., 2020). As previously reported, the mature differentiated octopaminergic neurons found in hemilineage B express the transcription factor Vestigial (Vg) (Landgraf et al., 2003). Here we find that a small number of immature postembryonic neurons (about 3-5) in close proximity to the MNB also express Vg (**Figure 7J.**). Within these immature neurons the expression of Vg and En are mutually exclusive. Using *Worniu-GAL4* to drive SR4VH we found that only the engrailed-negative cells are undergoing apoptosis (**Figure 7K**), i.e. the same small number of cells that express Vg. Their proximity to the MNB suggests that Vg-positive B cells (i.e. immature octopaminergic neurons) undergo an early death, very soon after they are born.

## DISCUSSION

### Midline neurons show hemilineage-based variations in different species

The midline neurons within the VNC of insects have long been a source of interest because they are homologous across species, yet show a diversity in cell numbers correlated with body form and function (Stevenson and Spörhase-Eichmann, 1995; Witten and Truman, 1998; Lacin et al., 2019). For example, flying insects have greater numbers of midline octopaminergic neurons within segments that control wings (Konings et al., 1988; Siegler et al., 1991, 2001; Eckert et al., 1992; Stevenson et al., 1992; Spörhase-Eichmann et al., 1992; Thompson and Siegler, 1993; Campbell et al., 1995; Monastirioti et al., 1995; Siegler and Pankhaniya, 1997; Jia and Siegler, 2002; Schlurmann and Hausen, 2003), while grasshoppers have more GABAergic neurons in the metathoracic/abdominal ganglia, which receives auditory input (Witten and Truman, 1998; Thompson and Siegler 1991). These two midline neuronal populations are derived from the same neuroblast, the MNB, which buds off multiple GMCs, each dividing once to generate a GABAergic (A cell) and an octopaminergic (B cell) neuron. Both A and B neurons are generated in equal numbers but, in all cases, the numbers of GABAergic (‘hemilineage A’) and octopaminergic (‘hemilineage B’) neurons within one segment are never the same. The greater numbers of GABAergic cells within each segment has been shown in grasshoppers (Jia and Siegler, 2002) and fruit flies (Truman et al., 2010) to be the result of removal by PCD of large numbers of cells from hemilineage B.

As suggested by our previous work, the ‘hemilineage’ emerges as a discrete developmental unit that shows common features of gene expression and function (Truman et al., 2010; Harris et al., 2015; Lacin and Truman, 2016; Lacin et al., 2019; Shepherd et al., 2019). Therefore, we refer here to the PCD found in the MNB lineage of grasshoppers as ‘hemilineage-specific PCD’. Following our observations of PCD during development in other insects, together with a vast body of knowledge on homology in insect nervous systems (Thomas et al., 1984; Kutsch and Breidbach, 1994), we suggest that variations in neural circuits between species is very likely set up by modifying hemilineages, with PCD playing a major role.

### Cell death during neurogenesis is widespread across insects

Our data show that PCD is extensive and widespread during neurogenesis in the CNS of insects, from the primitive firebrats *Thermobia domestica*, to true flies *Drosophila melanogaster* and the swift lousefly *Crataerina pallida*.

We wondered whether alterations in cell death may have contributed to adaptations in the VNC of flightless dipterans and found that a greater extent of PCD in the MNB lineage is responsible for abolishing the segment-specific difference in octopaminergic cell numbers in the swift lousefly *Crataerina pallida* – which has lost flight and gained adaptations to a parasitic lifestyle (Hagan, 1951; Bequaert, 1952; Hutson, 1984; Lehane, 2008; Walker and Rotherham, 2010b; Petersen et al., 2018). As octopaminergic neurons are involved in flight-related behaviours (Duch and Pflüger, 1999; Roeder, 2005; Brembs et al., 2007), we suggest that PCD has been co-opted in this lineage during the evolution of flightlessness in the swift louse. We believe that this PCD takes place early, in newly born neurons, as we have seen dying cells close to the MNB in all of our 20 pupae, from day 4 to day 23. Therefore, the decrease in octopaminergic cell number in the mesothoracic segment of swift louseflies is likely the result of increased hemilineage-specific PCD during evolution.

### Hemilineage-specific reductions in fiber diameter in the bee lousefly

In the wingless bee lousefly *Braula coeca,* we found clear reductions in the thickness of fibre tracts in several hemilineage bundles which in *Drosophila* are associated with flight-related behaviours (Harris et al., 2015). It remains for us to determine if early PCD takes place in these specific bee lice lineages during development and causes the reduction in bundle diameter that we see in flight hemilineages. As with most parasitic insects, bee and swift louseflies are impossible to maintain in the laboratory in the absence of their hosts and procuring them is a challenge (e.g. bee louseflies are now only found on two islands in the UK, while collecting swift louseflies is restricted to the summer months due to *Apus apus* migrations). Nonetheless, our observations that PCD is widespread across insects complements our findings in bee louseflies, strongly suggesting that an extensive PCD in flight hemilineages accompanied the loss of flight during evolution. Interestingly, the reduction in fibre diameter we see in bee louseflies was not evident in swift louseflies. This difference is likely due to the more significant changes to body plan in bee louseflies, i.e. a complete loss of the flight apparatus during evolution. Swift louseflies however still maintain vestigial wings and halteres (Walker and Rotherham, 2010), whereas bee louseflies have a severely reduced thorax, completely lacking wings, halteres and flight muscles (McAlister, 2018).

### Blocking death results in functional neurons that integrate into adult networks

Here we blocked PCD within the MNB lineage (called lineage 0 in the postembryonic literature) and found that ‘doomed’ neurons become octopaminergic, generate arborizations, and target the tectulum neuropil (Court et al., 2017). Previous work has shown that octopamine can induce and maintain rhythmic behaviours such as stepping movements and flight muscle contractions in locusts (Sombati and Hoyle, 1984) and walking, wing flicking and hindleg grooming in decapitated fruit flies (Yellman et al., 1997). Consistently, we find that our ‘undead’ hemilineage 0B cells can induce walking when activated thermogenetically in headless flies and show calcium activity during naturalistic bouts of locomotion. Thus, by blocking death, we have ‘resurrected’ functional neurons that are able to integrate into thoracic motor networks. Although this ‘dialling up’ of cell numbers is artificial in our system, it reveals how doomed neurons possess cryptic cellular phenotypes that can emerge when death is blocked, advocating for the evolvability of such a hemilineage based system.

While these undead hemilineage 0B octopaminergic neurons share many conserved features with wild-type cells, the variations we see in their morphology may act as a substrate for evolutionary change. A recent study has linked structural changes in the ventral nerve cord (VNC) with changes in behaviour between strains of *Drosophila melanogaster* (Mellert et al., 2016). Mellert *et al.,* show that hemilineage 12A in the mesothorax has variable bundle morphologies and that these correlate well with the time of flight initiation. Flight is used as an escape response and can be instrumental for predator evasion, one of the major evolutionary forces which have selected for flight in insects in the first place (Dudley, 2002). With this in mind, it seems plausible that changes in either neuron number and/or innovations in ‘undead’ neuron structure could affect adult behaviour and be ultimately adaptive. Recently it has been shown that undead sensory neurons that are functional, integrate, and appear to be tuned to specific odours (Prieto-Godino et al., 2020). Importantly, Prieto-Godino *et al*. also show that there is cell number variation in this neuronal population across drosophilids and that blocking PCD in *melanogaster* results in the survival of mosquito-like CO_2_-sensing neurons in the maxillary palps. This suggests that both the central and peripheral nervous system may use similar modes of early PCD to sculpt circuits during the evolution of true flies.

### Hemilineage-specific cell death occurs in newly born neurons

To help us understand more about the patterning of PCD we designed and used a new effector caspase probe SR4VH allowed us to interrogate the extent and dynamics of hemilineage-specific cell death. It shows us that an early-onset PCD is responsible for the elimination of postembryonic neurons in the fly VNC, that this happens throughout the entire 3.5 days of postembryonic neurogenesis, and is hemilineage-specific. Although PCD has been reported as a fate within lineages in the embryo (Karcavich and Doe, 2005; Rogulja-Ortmann et al., 2007), the impact of this ‘early’ and hemilineage-specific PCD on the construction of the adult network has yet to be fully appreciated. This type of PCD is responsible for removing almost half of all postembryonic neurons that are born in the fly (Truman et al., 2010). Until now, the most frequently reported type of neuronal death described in insects has been the hormonally-regulated PCD that removes mature neurons during the narrow developmental windows at the beginning of metamorphosis and within the first day after adult eclosion (Draizen et al., 1999; Lee et al., 2013). We now know from our data that these make up only a small fraction, compared to the total number of neuronal deaths in the fly.

Our SR4VH probe allows us to see that newly born neurons initiate cell death within the first 5.5 hours after birth. We can capture different stages of cell death: with young cells at very initial stages of PCD located closer to the neuroblast, while older cells at more advanced stages of PCD with RFP labelled cell membranes found close to the lineage bundle.

Importantly, death happens before neurites have extended, strongly suggesting that this PCD is not an analogue of neurotrophic death, found in vertebrates - where neuron-target interactions play a major role in the decisions of cell survival (Dekkers and Barde, 2013). This the dynamics of hemilineage-specific cell death we see with SR4VH fits well with the patterns that we find in *Thermobia domestica* and *Crataerina pallida* suggesting that they too undergo a death with rapid onset following division.

The critical question that this data brings into focus is how early-onset PCD is orchestrated, especially that an early intrinsically-determined mode of cell death seems to be widespread across animals, from *C. elegans* to mice (Southwell et al., 2012; Fricker et al., 2018). Early PCD likely involves a combination of intrinsic patterning and cell-cell interactions between sibling neurons which ultimately deploy the activity of proapoptotic genes *rpr*, *hid*, *grim* and/or *skl*. Previously, patterning genes such as *Ubx* have been shown to contribute to the survival of hemilineages in the thoracic VNC in a parasegment-specific manner (Marin et al., 2012), while the transcription factor Unc-4 has been recently demonstrated to provide neural identity to hemilineages involved in flight (Lacin et al., 2020). Such spatial patterning is also required to establish NB identity which, in turn, can determine which hemilineage is maintained and which dies. In the developing optic lobe Bertet *et al*. have shown that the temporal sequence of transcription factors expressed in neuroblasts (inherited by the GMC and newly-born neurons) produces a switch in the selective survival of one hemilineage over the other (Bertet et al., 2014). In this manner, the changes to NB identity which have been documented in other insects (Biffar and Stollewerk, 2014), despite a conserved NB array and progeny, could in turn influence the pattern of PCD. Alongside, cell-cell interactions between newly-born neurons could influence fate choices. The requirement of interactions between newly-born siblings in determining asymmetric fates has been shown in the grasshopper VNC (Doe et al., 1985; Kuwada and Goodman, 1985), although this exact same mechanism has not been demonstrated in *Drosophila*.

The spatio-temporal pattern of death revealed with SR4VH shows us that understanding the molecular control of hemilineage-based death is the key question going forward and is likely to provide insight into how networks evolve.

## Conclusions

Here we have shown that undead neurons elaborate complex arborizations, express distinct transmitter identities and function. We find hemilineage-based, ‘early’ programmed cell death is widespread during the development of the CNS of insects from the primitive Firebrats, to most derived true flies. Early cell death appears to be a specific subtype of PCD present across animals. Understanding how early PCD is specified across species should help us elucidate how nervous systems are built and evolve. Our exploration of homologous lineages in flightless dipterans shows that changes in the extent and pattern of PCD are evident with changes in body plan. As the evolutionary changes seen in neural networks ultimately result from heritable differences in developmental processes, our future endeavours will be directed towards elucidating how genetic programs are deployed to establish the pattern of PCD. The cellular leitmotif of hemilineage-based cell death, we present here, provides us with something tangible that we can search for. Thus, we suggest that viewing the evolution of insect nervous systems through the lens of the ‘hemilineage’ will be critical for understanding how development brings about adaptive changes in neural network motifs.

## SUPPLEMENTARY FIGURES

**Figure S1.**
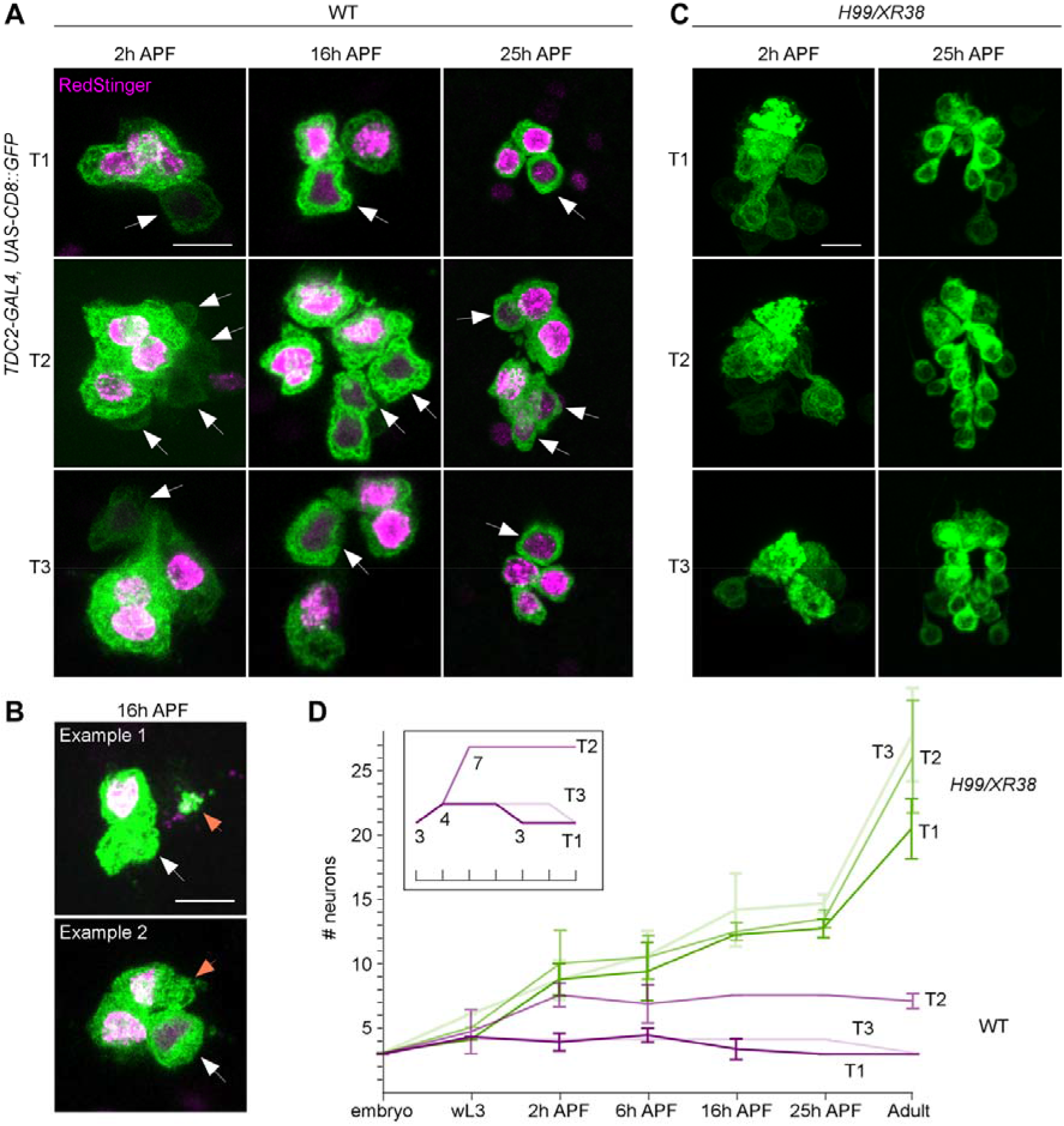
Postembryonic development of lineage 0B in wild-type and *H99/XR38* flies. **(A)** CD8::GFP expression driven by *TDC2-GAL4* in octopaminergic neurons from hemilineage 0B in the VNC of wild-type flies in T1 (1^st^ row), T2 (2^nd^ row) and T3 (3^rd^ row) during pupal development at 2 hours after puparium formation (2h APF, 1^st^ column), 16h APF (2^nd^ column) and 25 hAPF (3^rd^ column). One embryonically-born cell is replaced during postembryonic neurogenesis in T1 and T2, while roughly 4 postembryonic octopaminergic neurons are added in T2. White arrows indicate newly-born neurons with faint GFP and no RedStinger visible at 2hAPF (arrows). RedStinger fluorescence becomes visible at 16-25hAPF, as the protein matures (arrows). Scale bar, 10 μm. (**B**) Accumulations of GFP and RedStinger (orange arrow) in the proximity of the prothoracic cell cluster, alongside the presence of one cell with dim RedStinger fluorescence (white arrow) suggest one embryonic cell has died at 16hAPF. Scale bar, 10 μm. (**C**) CD8::GFP expression driven by *TDC2-GAL4* in octopaminergic neurons from hemilineage 0B in the VNC of *H99/XR38* flies in T1 (1^st^ row), T2 (2^nd^ row) and T3 (3^rd^ row) during pupal development at 2h APF (1^st^ column) and 25h APF (2^nd^ column). Not all undead neurons present in the adult have matured by 25h APF. Scale bar, 10 μm. (**D**) Graphical representation of the number of *TDC2-GAL4*-expressing neurons throughout development in wild-type (magenta) and *H99/XR38* mutant flies (green), where X represents time and Y represents the number of cells. Individual lines in the graph depict cell numbers in different thoracic segments. Error bars represent standard deviation. The inset represents a compressed idealised graphical representation for wild-type flies; marks on the X axis are the same; numbers at branch points represent cell numbers. In wild-type flies both T1 and T3 acquire an additional cell at 2h APF and lose one by the end of metamorphosis (magenta). Roughly half of undead neurons are present at 25 hAPF, with the other half maturing by the end of metamorphosis (green).

### Comment on S1 data

#### The implication of additional postembryonic octopaminergic neurons born in the in the wild-type larva

It is well known that there are 3-4 extra octopaminergic neurons generated postembryonically in the wing bearing second (T2) thoracic neuromere in adult flies compared to the 3 cells found in each thoracic segment in larvae [29]. Until our inquiry into the timeline of postembryonic neurogenesis, it was thought that all octopaminergic neurons born in the embryo persist into adulthood, but to our surprise, that is not the case. We noticed that one postembryonic octopaminergic cell is born in wild-type flies in both the first (T1) and third (T3) thoracic segments, bringing the total number of cells to 4 (**Figure S1A,D**). Instead, the total number of 3 cells per cluster in T1 and T3 in adult flies is achieved via *hormonally gated metamorphic cell death of an embryonically born mature neuron* at larval-pupal stages (**Figure S1A,B,D**). We have shown here for the first time that wild-type octopaminergic neurons are produced in lineage 0 in all three thoracic neuromeres during postembryonic neurogenesis. This fits well with our observations that some *Dronc*-null singleton clones generated using MARCM showed bilateral symmetry. As the majority of these were in the mesothorax (see also **Figure S2A,D**), where it is more likely to label wild-type neurons purely by chance (with up to 4 postembryonic neurons produced here and only 1 each in T1 and T3), we think these may be indeed wild-type. Furthermore, we only ever encountered mixed MARCM clones containing one bilaterally-symmetric and several turning neurites in the mesothorax (see also **Figure S2A,D**). For behavioural experiments we excluded all MARCM *Dronc* clones with bifurcating neurons to ensure that we only interpreted the activity of undead neurons.

**Figure S2.**
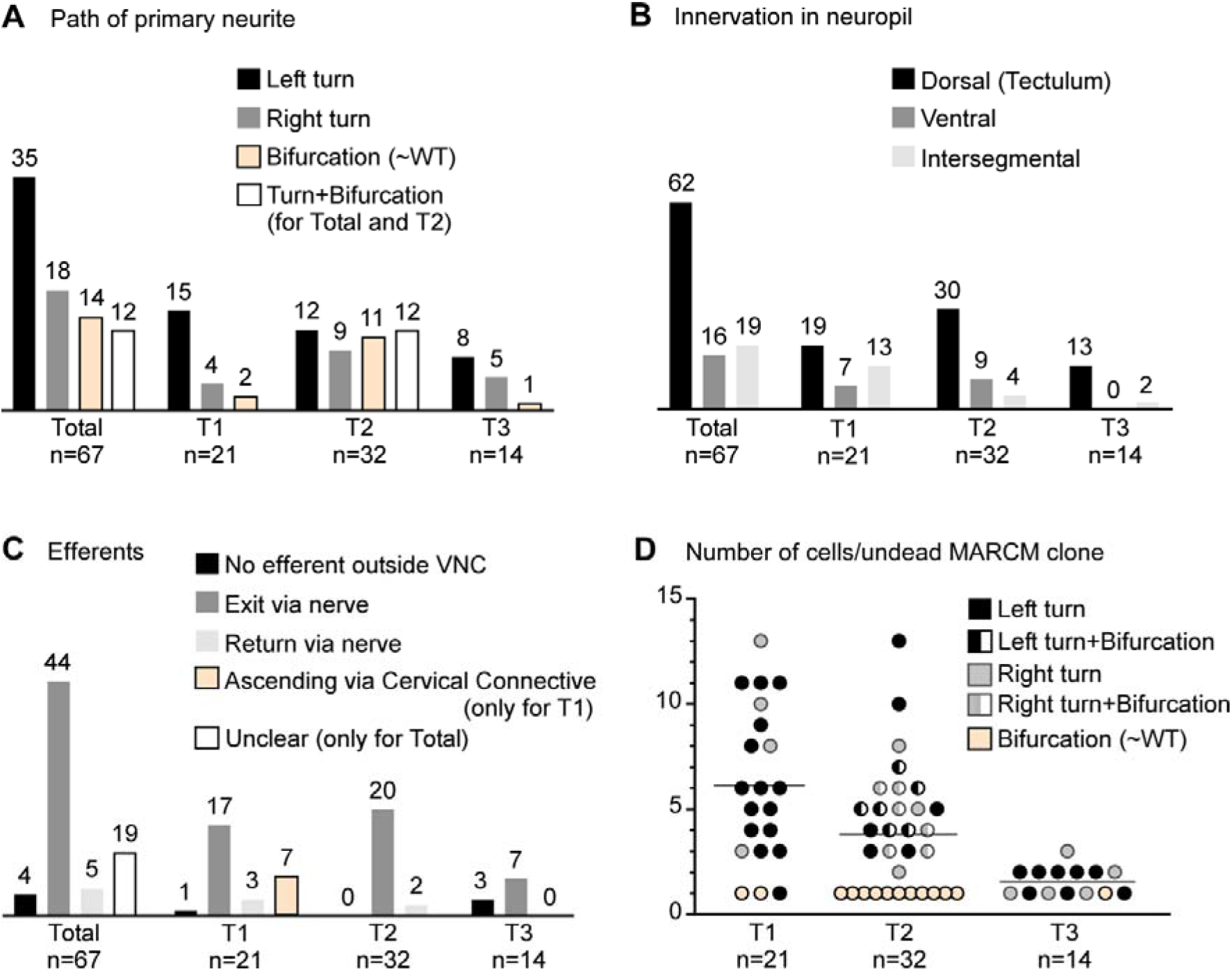
Quantification of undead neuron morphology. **(A)** Quantification of the primary neurite path. Most undead neurons arborize to the left. **(B)** Quantification of neuropil innervation. Most undead neurons branched extensively in the dorsal neuropil. Undead neurons in T3 never innervated the ventral neuropil. Few undead neurons from each thoracic segment projected neurites into adjacent segments. **(C)** Quantification of efferent neurites. Most undead neurons joined a thoracic nerve heading towards the periphery. Some undead neurons in T1 sent branches through the cervical connective. (**D**) Quantification of the number of undead neurons in MARCM clones for each thoracic segment color-coded by primary neurite path. The number of undead neurons varied across MARCM clones, with up to 13 in T1 and T2 and up to 3 in T3. Most single-celled MARCM clones contained undead neurons with a wild-type morphology (beige circles) and most undead neurons with a wild-type morphology were encountered in T2. Mixed clones (half-filled circles) made up of one bifurcating neuron and several neurons turning either left or right were encountered only in T2.

**Figure S3.**
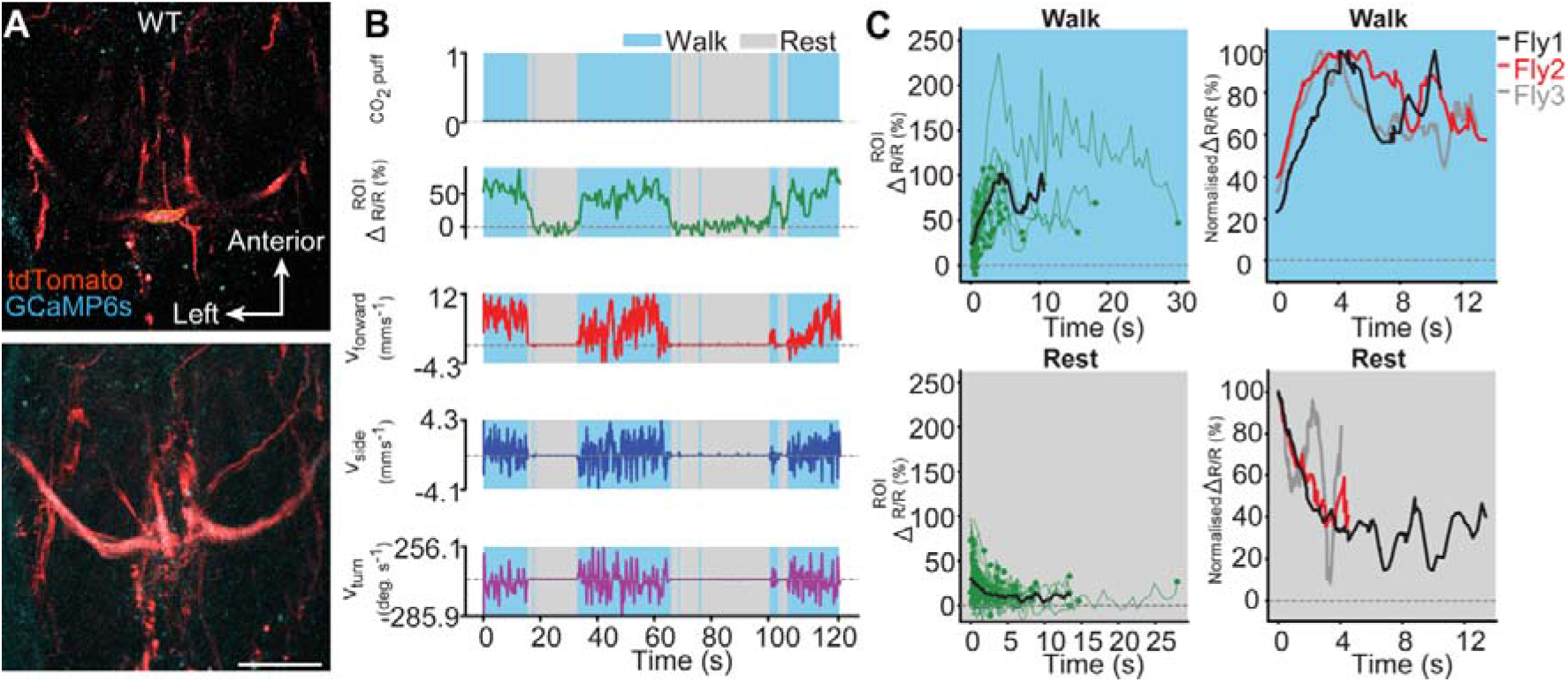
Wild-type octopaminergic neurons are active during walking in intact adult *Drosophila*. **(A)** Raw 2-photon image of *TDC2-GAL4-positive* neurons co-expressing tdTomato (red) and GCaMP6s (cyan) in wild-type flies (**top**). Region-of-interest used to calculate %ΔR/R is outlined (yellow). Standard deviation z-projection of a dorsal-ventral image stack of the functional imaging region-of-interest in **b** (**bottom**). Scale bar, 50 μm. **(B)** Representative behavioural and functional imaging data in wild-type flies. Shown are: CO_2_ stimulation (black, no stimulation), %ΔR/R (ratio of GCaMP6s / tdTomato) signal (green), and ball rotations indicating forward walking (red), sideways walking (blue), and turning (purple). The behaviour of the fly was classified as either walking (light blue), or resting (grey) by applying a threshold on ball rotation speed. **(C) (left)** Individual (green) and average (black) %ΔR/R traces within each behavioural epoch for walking (n = 77) and resting (n = 80) events processed from 720 s of imaging data. Solid green circles indicate the end of a behavioural epoch. The average trace (black line) was calculated for only periods with 4 or more traces. (**right**) Normalised average %ΔR/R traces for three different flies during walking and resting. The average (black) trace is the same as in the left panel.

**Figure S4.**
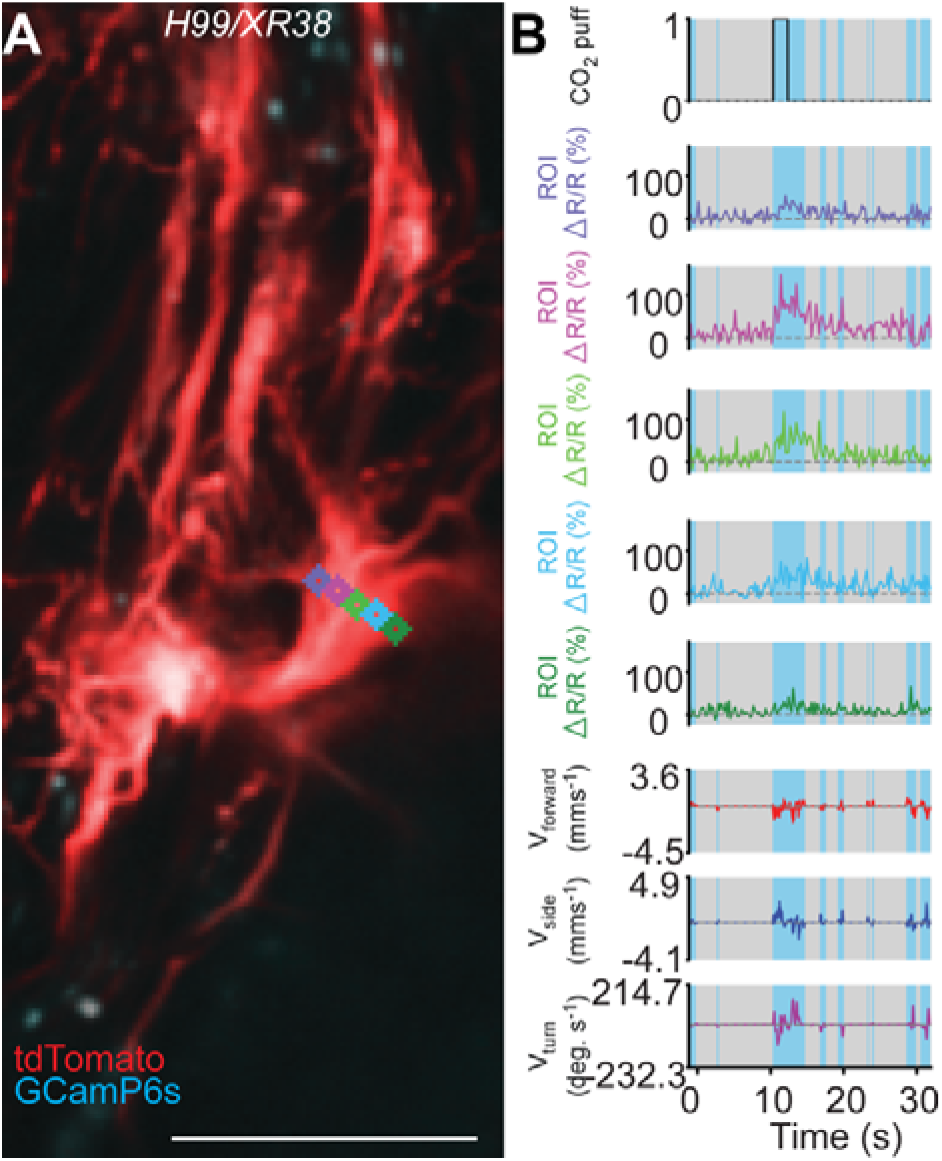
Subregion analysis of calcium signals along the width of a primary neurite in a *H99/XR38* animal. **(A)** Time-averaged projection of the imaging plane for *TDC2-GAL4*-positive neurons co-expressing tdTomato (red) and GCaMP6s (cyan) in a *H99/XR38* animal. Regions-of-interest used to calculate %ΔR/R (color-coded) are overlaid on top of time-projected and optic flow registered 2-photon images. Scale bar, 50 μm. **(B)** Representative behavioural and functional imaging data for this animal. Shown are: CO_2_ stimulation (black), %ΔR/R (ratio of GCaMP6s / tdTomato) signal (color-coded for each ROI as in panel **a**), and ball rotations indicating forward walking (red), sideways walking (blue), and turning (purple). The behaviour of the fly was classified as either walking (light blue), or resting (grey) by applying a threshold on ball rotation speed.

**Figure S5.**
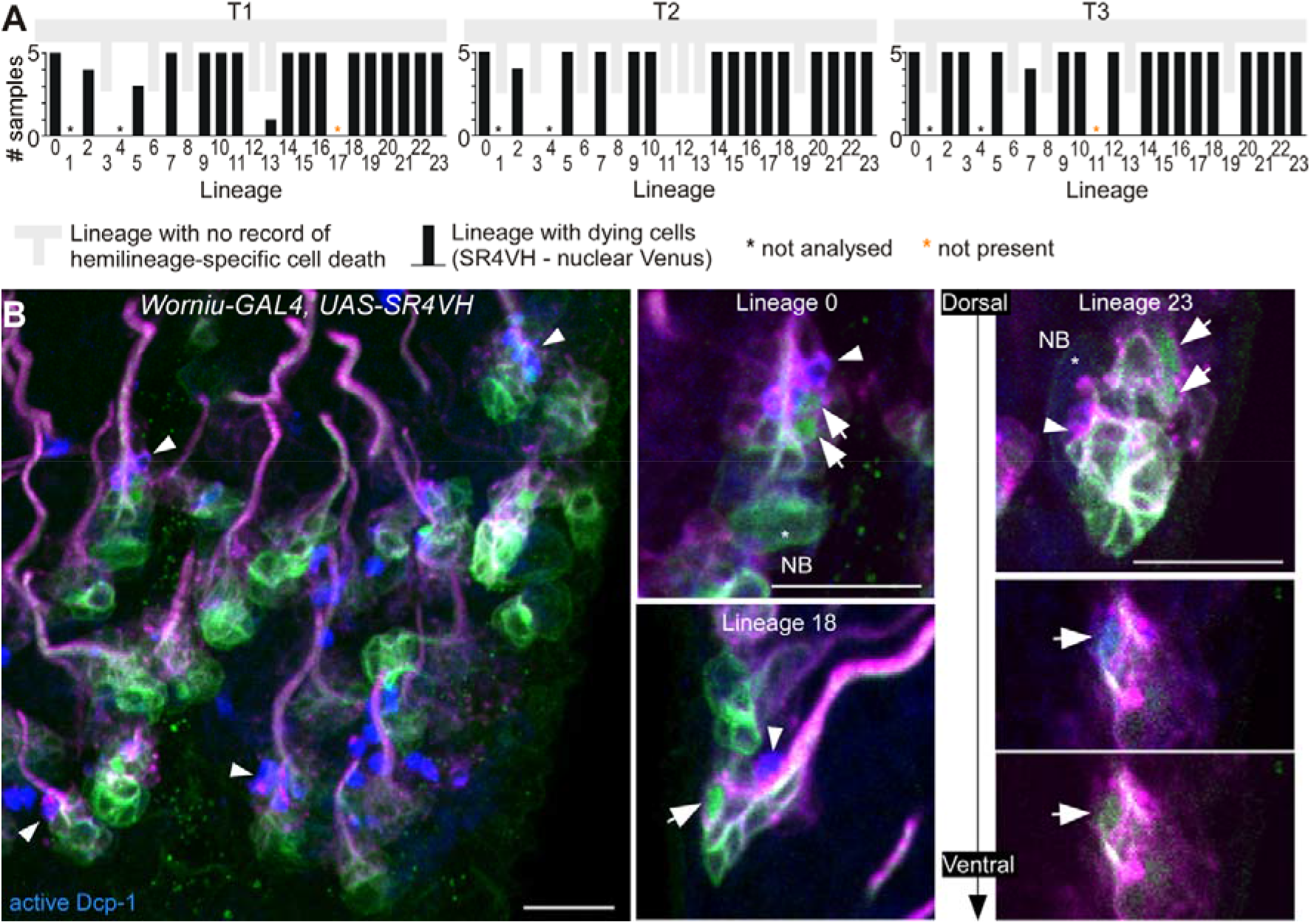
SR4VH reveals successive stages of cell death in lineages with doomed hemilineages. **(A)** Quantification of lineages with dying cells as reported with SR4VH in each thoracic segment from 5 wandering 3^rd^ instar larvae. Back bars represent the number of samples in which dying cells were encountered in at least one hemisegment for that lineage. Grey bars represent lineages with no record of hemilineage-specific cell death as reported in Truman et al., 2010. **(B)** Antibodies for active Dcp-1 label cells located close to the lineage bundle in doomed lineages from larvae expressing SR4VH driven by *Worniu-GAL4*. Right panels show examples of doomed lineages 0, 18 and 23 containing cells at subsequent stages of cell death. White arrows indicate: nuclear Venus without cleaved Dcp-1 (early-stage), Venus and Dcp-1 colocalization (mid-stage), and pyknotic cells/dead cell membranes with RFP and cleaved Dcp-1 (late-stage). Scale bars, 10 μm. *n* = 7.

## SUPPLEMENTARY VIDEOS

### Video S1

Video recordings of control and ‘undead’ decapitated flies during thermogenetic activation. Examples from each behavioural category showing responses to heat-activation in negative control (top left), positive control (top right), MARCM control (bottom left) and undead neurons (bottom right). MARCM control and undead neuron animals were used for extracting the centroid trace provided in **Fig. 3, B** **and** **C**. The increase in temperature is displayed in the bottom right corner. Frames represent recordings from 30 to 70s.

### Video S2

Recording of 2-photon calcium imaging in undead neurons. Synchronized front and side camera behaviour videography (**bottom-right**), and 2-photon imaging data (**top-right**) used for the data analysis provided in **Fig. 3, H-J**.

### Video S3

Z-stack of the imaging area for GCaMP6s activity in undead neurons. A videography showing the imaging plane at different depths of the prothoracic segment corresponding to **Figure 3H.** The thicker left branch likely includes mostly undead neurons.

### Video S4

Subregion neuronal activity patterns during walking. Imaging data used for analysis in **Fig. S4**. Shown are %ΔR/R traces for R

## MATERIALS AND METHODS

### Animals

We used the following *Drosophila melanogaster* stocks: *Worniu-GAL4; Dr/TM3, Ubx-LacZ, Sb* (BL 56553), *Tdc2-GAL4* (BL 9313), *OK371-GAL4* (BL 26160), *UAS-SR4VH* (described here), *UAS-CD8::GFP* (BL 5137), *UAS-tdTomato-p2A-GCaMP6s* (*28*) (kind gift from M. Dickinson), *H99/TM3, Sb* (BL 1576), *XR38/TM3*, Sb, *If/CyO; dronc*^Δ*A8*^, *FRT2A/TM6*^β^*, Tb, Hu* (*37*) and *hs-flp;; TubP-GAL80, FRT2A/TM3, Sb* (*16*).

Firebrat adults of *Thermobia domestica* were obtained from Buzzard Reptile and Aquatics (buzzardreptile.co.uk) and reared on a diet of fish flakes and wholemeal bran at 40°C in darkness inside a humid plastic container. A staging series was calculated by time to hatching.

*Crataerina pallida* swift lousefly adults were collected from swift (*Apus apus*) nesting boxes fitted behind the louvres of belfry windows from churches in Cambridgeshire and Suffolk (UK) with the help of local conservationists Simon Evans, Richard Newell and Bill Murrells. Swift louseflies were kept at 20°C on a 12h dark:12h light cycle until dissected. Pregnant females, recognised by their enlarged and translucent abdomen through which larvae or prepupae could be detected, were kept separately and checked daily for pupa ejection. The day in which a pupa was laid was defined as ‘Day 0’ of external development (outside the mother’s abdomen).

*Braula coeca* bee lousefly adults were obtained from a black bee (*Apis mellifera mellifera*) colony on the Isle of Colonsay, UK (kind gift from A. Abrahams). Bee louseflies were shipped by post in small cages containing worker bees feeding on bee fondant. The black bees and bee louseflies were anesthetized by placing the cage on a CO_2_ pad and the bee louseflies were removed for dissection.

### Construction of *UAS-SR4VH*

SR4VH was constructed by standard molecular biology procedures. It comprises the myristylation signal of *Drosophila* Src64B (amino acids 1-95), a monomeric red fluorescent protein mRFP1 (*38*), a linker that contains four DEVD sites, a yellow florescent protein Venus (*39*), and a nuclear localization signal of *Drosophila* histone H2B (amino acids 1-51). While the design is similar to the previously reported caspase probe Apoliner (*40*), the Src64B myristylation signal and the H2B NLS offers better membrane and nuclear localization, respectively, and four DEVD sites are expected to provide higher sensitivity. The probe was cloned in pUAST (*41*) and introduced into the *Drosophila* genome by P element-mediated transformation.

### Immunohistochemistry and chemical staining

*Drosophila* larvae were dissected in PBS without anaesthesia. Firebrat embryos were removed from their chorion and dissected using minuten pins. *Drosophila*, swift lousefly and bee lousefly adults were anesthetised on ice, briefly submerged in absolute ethanol and dissected in PBS. Swift lousefly pupae were immobilised on double sided sticky tape, removed from their pupal case using forceps and dissected in PBS without anaesthesia. Samples were fixed in 3.6% paraformaldehyde in PBS for 30 min (larvae and pupae) or 1h (adults), washed 3 times in 0.3% PBST (0.3% Triton-X100 in PBS, Sigma-Aldrich), blocked in 5% goat serum (Sigma-Aldrich) in PBST for 1h and incubated with primary antibodies in block for 1-3 days at 4°C (*Drosophila*, bee louseflies, swift lousefly pupae), room temperature (firebrats) or 37°C to increase antibody penetration (swift lousefly adults; block supplemented with 0.02% NaN_3_ to prevent microbial growth). Samples were then washed 4 times throughout the day in PBST and incubated with secondary antibodies in block for a further 1-3 days, followed by final washes in PBST and PBS. Brains and VNCs were mounted on poly-L-lysine-coated coverslip, dehydrated in increasing serial concentrations of ethanol (15%, 30%, 70%, 80%, 90% and twice in 100%) for 5min each, dipped once in xylene, then incubated twice for 5min in fresh xylene. A droplet of DePeX (EMS) was added on top of the mounted sample and the coverslip was placed face-down on a glass slide.

We used the following primary antibodies: chicken anti-GFP (1:500; ab13970, Abcam), mouse anti-Neuroglian (1:50; BP 104, Developmental Studies Hybridoma Bank), rabbit anti-cleaved *Drosophila* Dcp1 (1:100; Asp216, Cell Signaling), guinea pig anti-Syncrip (1:100; kind gift from I. Davis; to label neuroblasts and early progeny in lineages - Jim Truman, personal communication), mouse anti-Engrailed/Invected (1:2; 4D9, Developmental Studies Hybridoma Bank), rabbit anti-DVGLUT C-terminus (*43*) (1:5000; AB_2490071, kind gift from H. Aberle), rat anti-tyramine β-hydroxylase (*30*) (1:50; TBH, kind gift from M. Monastirioti), rabbit anti-vsg (1:400; kind gift from Sean Carroll and Kirsten Gruss) and rabbit anti-GABA (1:100; AB_572234, Imunostar).

Secondary antibodies were Alexa Fluor® 488-conjugated goat anti-chicken (1:500; A11039, Invitrogen, Thermo Fisher Scientific), Alexa Fluor® 488-conjugated goat anti-rabbit (1:500; A11070, Invitrogen, Thermo Fisher Scientific), Cy™ 3-conjugated donkey anti-rabbit (1:500; 711-006-152, Jackson ImmunoResearch), Cy™ 5-conjugated donkey anti-mouse (1:500; 715-006-151, Jackson ImmunoResearch), Alexa Fluor® 488-conjugated donkey anti-rat cross-adsorbed against mouse (1:100; 712-545-153, Jackson ImmunoResearch), Alexa Fluor® 488-conjugated donkey anti-guinea pig (1:500; A11073, Invitrogen, Thermo Fisher Scientific).

In firebrat embryos we detected dying cells using the Click-iT Plus TUNEL assay kit (C10618, *life* technologies). To stain cell nuclei and neuropil, firebrat samples were incubated with DAPI (1:10000; D9542, Sigma-Aldrich) and Phalloidin-488 (1:100, *life* technologies) in PBST for 30 min at room temperature. Incubations were carried out following secondary antibody treatment. Samples were then washed in PBST and PBS, and mounted.

### EdU treatment

To label proliferating cells and their progeny we used the Click-iT EdU imaging Kit (C10337, *life* technologies). Freshly dissected nervous systems from swift lousefly pupae were incubated in EdU 1:1000 in PBS at room temperature for 1-3h on a shaker, rinsed with PBS and fixed in cold buffered formaldehyde 3.6% in PBS for 30min. Samples were then stained using the immunohistochemistry protocol described above. The colour reaction for EdU was carried out as instructed by the vendor after the secondary antibodies were washed out.

### Generation of Undead neuron MARCM clones

To induce mitotic clones of undead neurons, rescued from programmed cell death, we used the mosaic analysis with a repressible cell marker technique (*43*). 0-4h first instar larvae resulting from crossing females of the genotype *hs-flp; ; TubP-GAL80, FRT2A/TM3, Sb* with; *TDC2-GAL4, UAS-CD8::GFP, UAS-TrpA1; dronc*^Δ*A8*^, *FRT2A/TM6β, Tb, Hu* males were heat-shocked at 37°C in a plastic food vial placed in a water bath for either 1h or 45min, followed by 45min at room temperature and a second incubation period at 37°C for 30min. After heat-shock, larvae were immediately returned to 23 or 25°C. Cell death was blocked in clones homozygous for the loss-of-function allele of the initiator caspase dronc. Because we used the octopaminergic driver line *TDC2-GAL4* to induce the expression of CD8::GFP and TrpA1, we were able to visualise and thermogenetically activate only postembryonic neurons of hemilineage 0B. A small number of wild-type octopaminergic neurons are born during postembryonic neurogenesis (1 in T1 and T3, 4-5 in T2, data now shown). To ensure the characterisation of undead neurons only, MARCM clones including a bilaterally symmetrical primary neurite were excluded from analysis.

### Thermogenetic activation and video recordings

Prior to recordings, 2-6-day-old males of *Drosophila melanogaster* (*hs-flp/+; Tdc2-GAL4, UAS-CD8::GFP, UAS-dTRPA1/+; dronc*^Δ*A8*^, FRT2A/TubP-GAL80, FRT2A) reared at 23 or 25°C in a 12h:12h light:dark cycle were anesthetised on ice and decapitated using a pair of micro spring scissors in under 3min. We used males as we found they are more responsive to octopamine release by thermogenetic activation than females (data not shown). The headless flies were brushed back into a food vial placed on its side and left to recover for at least 1 hour. To generate the heat ramp required to thermogenetically activate undead neurons, we used a 12V thermoelectric Peltier plate (model: TEC1-12706, size: 40mm x 40mm x 3.6mm) connected to a DC power supply (HY3005D, Rapid Electronics) set at a constant current of 0.46A, with a variable voltage, calibrated using an infrared laser thermometer (N92FX, Maplin). These settings generated a temperature ramp which lasted 70s from 22°C to 34°C. Videos were recorded at 25 fps using a Sony NEX-5N digital camera (kindly provided by Ian Wynne) mounted to a stereo microscope. A piece of graph paper was used for spatial calibration. To match the presence of undead neurons with behaviour, each decapitated fly used for thermogenetic activation was indexed and prepared for dissection and immunostaining.

### Two-photon calcium imaging in behaving intact flies

The method for *in vivo* two-photon imaging of the VNC in behaving adult *Drosophila* is described in (*28*). Briefly, flies were anesthetized through cooling and then mounted onto custom imaging stages. The dorsal thoracic cuticle was removed and indirect flight muscles were left to degrade over the course of 1h. Subsequently, the proventriculus and salivary glands were resected to gain optical access to the VNC.

Horizontal sections of the T1 leg ganglion were imaged using galvo-galvo scanning. For control animals, the bifurcation point of TDC-positive neurites were imaged to circumvent ROI disappearances caused by movement. For animals harbouring undead *TDC2-GAL4*-positive neurons, the thickest branch of the axonal bifurcation was chosen because they were most likely to contain undead neurites. Image dimensions ranged between 512×512 and 320×320, resulting in 1.6 to 3.4 fps data acquisition. Imaging areas ranged between 92×92 μm and 149×149 μm. Laser power was held at ~8 mW.

### Data analysis for 2-photon imaging in behaving *Drosophila*

Python scripts (modified from (*28*)) were used to extract ROI fluorescence traces and to compute spherical treadmill ball rotations. Walking epochs were determined by placing a threshold on ball rotations, which were first converted into anterior-posterior (v_forward_) and medial-lateral (v_side_) speeds (1 rot s^−1^ = 31.42 mm s^−1^) and into degrees s^−1^ (1 rot s^−1^◻=◻360° s^−1^) for yaw (v_rotation_) movements. Thresholds were 0.12 mm, 0.12 mm, and 5 degrees, respectively. Periods below these thresholds were considered ‘resting’ while other periods were considered ‘walking’. Fluorescence traces for epochs with the same behaviour were aligned by start point to compute average %ΔR/R traces for specific actions.

To calculate fluorescence traces for small subregions-of-interest across neuritic bundles containing both undead and wild-type neurites, images were registered using an optic flow method described in (*28*). This registration served to minimize motion artefacts. Analysis was limited to a period with no warping artefacts and no ROI disappearance. Subregions were manually selected as small circular ROIs across the neuritic bundle of the registered image. Fluorescence values were then computed from each sub-ROI.

### Confocal imaging and image processing

Images were acquired using a Zeiss LSM 510 or a Zeiss LSM 800 confocal microscope at a magnification of 20x or 40x with optical sections taken at 1μm intervals. The resulting images were examined and processed using Fiji (https://imagej.net/Fiji). Some images were manually cropped using the Freehand Selection tool to remove debris or to cut out neuronal lineages in *; Worniu-GAL4, UAS-SR4VH;* samples.

### Fluorescence intensity plots

To generate fluorescence intensity along Line plots, we used the Plot Profile tool in Fiji to extract raw fluorescence intensity values for the RFP and Venus channels. The values were imported into MATLAB (R2018a, MathWorks) and normalised by dividing all fluorescence intensity values to the maximum value encountered along each Line. In this manner, all fluorescence intensity along Line plots have a common scale from 0 to 1, with 1 being the highest value encountered along that Line.

### Analysis of thermogenetic activation

Decapitated flies were considered to be walking if they covered a distance greater than 1 body length and moved their legs in a coordinated sequence from T3 to T2 to T1 at least once on each side (11). Forward, backward and sideway movements were all interpreted as walking when both aforementioned conditions were respected. To generate fly body traces video recordings were imported in MATLAB (R2018a, MathWorks) and the centroid of the decapitated fly (located on the scutellum) was extracted from each frame using a custom-written script. Each frame was converted into a grayscale image, its contrast enhanced using contrast-limited adaptive histogram equalization, filtered using a gaussian smoothing kernel with a standard deviation of 4, binarized using a custom threshold and the geometric centre of the fly body automatically extracted and stored in an array. To confirm that the centroid detection was accurate, a red dot with the centroid coordinates was superimposed onto each frame of the original recording and the annotated movie was saved for manual inspection.

### Quantification of 3A/3B bundle diameters

For calculating 3A/3B hemilineage bundle diameter ratios in fruit flies and bee louseflies, we generated transverse rendered maximum intensity projections of inverted grayscale confocal stacks for the pro- and mesothorax (T1 and T2) and frontal projections for the metathorax (T3). Optical sections were selected to include the common lineage bundle and the individual hemilineage bundles after their split. Diameter measurements were taken at the widest point within 5 μm of the bundle split using the Straight Line tool in Fiji and ratios were calculated by dividing the diameter of hemilineage 3A to that of 3B.

### Statistical analysis

For comparing neuron numbers, 3A/3B bundle diameter and T2/T1 number of neurons, data was tested for normal distribution using the Kolmogorov-Smirnov test and visualisation of Normal Q-Q plots. Differences between groups were analysed using either the independent samples t-test for normally distributed data, Welch’s test if data failed to meet the homogeneity of variances assumption or Mann-Whitney t-tests if data failed to meet the normality and homogeneity of variances assumptions of the independent samples t-test.

For comparing the number of flies which walked in each experimental group, we performed a Pearson chi-squared test and interpreted the resulting exact significance if the minimum expected count was greater than 5, or the Fisher’s Exact Test 2-sided significance if the minimum expected count was lower than 5 in at least one cell of the contingency table. To correct for multiple comparisons we performed a Bonferroni correction (i.e. p values were multiplied by 6, the total number of pairwise tests).

All statistical tests were performed in SPSS Statistics 23 (IBM) with an α set at 0.05. In all figures, bars represent means ± standard deviation; *P\** < 0.05, *P\*\*\** < 0.001, *P^ns^* = not significant.

## ACKNOWLEDGEMENTS

We would like to thank Richard Benton and Lucia Prieto-Godino for discussions and sharing data. We thank Kristin White and Bloomington Drosophila Stock Center (NIH P40OD018537) for sharing flies; Maria Monastirioti, Hermann Aberle, Ilan Davis, Kirsten Gruss, Sean Carroll and Developmental Studies Hybridoma Bank (NICHD of the NIH, University of Iowa) for antibodies. We are indebted to Simon Evans, Richard Newell and Bill Murrells for their kind help in collecting swift lice and Andrew Abrams for sending us bee lice. We are grateful to Andrew M. Dacks and Hans-Joachim Pflüger for providing unpublished data on *Manduca* octopaminergic neurons. We would like to thank Ian Wynne for his camera. We also thank Matthias Landgraf, David Shepherd, Jon Clarke and Sanjay Sane for reading the manuscript. Funding: Williams: BBSRC BB/P025552/1 and BB/L022672/1. Ramdya: SNSF Project Grant: 175667; Eccellenza Grant: 181239; R’Equip Grant: 177102.

## AUTHOR CONTRIBUTIONS

D.W. and S.P. conceived the study and designed the experiments, apart from *in vivo* imaging; collected *C.pallida* in the field, and wrote the first draft of the manuscript. S.P. performed the experiments apart from *in vivo* imaging. P.R. and C.C. conceived and performed the *in vivo* imaging experiment. S.K. designed and made the genetically encoded caspase reporter SR4VH. C.S. generated the *T.domestica* data. P.R. and S.K. edited the final manuscript drafts.

## REFERENCES

Allen, A.M., Neville, M.C., Birtles, S., Croset, V., Treiber, C.D., Waddell, S., Goodwin, S.F., 2020. A single-cell transcriptomic atlas of the adult Drosophila ventral nerve cord. Elife 9. doi:10.7554/eLife.54074

Bardet, P.-L., Kolahgar, G., Mynett, A., Miguel-Aliaga, I., Briscoe, J., Meier, P., Vincent, J.-P., 2008. A fluorescent reporter of caspase activity for live imaging. Proc. Natl. Acad. Sci. USA 105, 13901–13905. doi:10.1073/pnas.0806983105

Bate, M.C., 1976. Embryogenesis of an insect nervous system I. A map of the thoracic and abdominal neuroblasts in Locusta migratoria. Journal of Embryology and Experimental Morphology 35, 107–123.

Bequaert, J.C., 1952. The Hippoboscidae or Louse-Flies (Díptera) of Mammals and Birds. Part I. Structure, Physiology and Natural History. Entomologica Americana 32, 1–209.

Bertet, C., Li, X., Erclik, T., Cavey, M., Wells, B., Desplan, C., 2014. Temporal patterning of neuroblasts controls Notch-mediated cell survival through regulation of Hid or Reaper. Cell 158, 1173–1186. doi:10.1016/j.cell.2014.07.045

Biffar, L., Stollewerk, A., 2014. Conservation and evolutionary modifications of neuroblast expression patterns in insects. Dev. Biol. 388, 103–116. doi:10.1016/j.ydbio.2014.01.028

Booker, R., Truman, J.W., 1987. Postembryonic neurogenesis in the CNS of the tobacco hornworm, Manduca sexta. I. Neuroblast arrays and the fate of their progeny during metamorphosis. J. Comp. Neurol. 255, 548–559. doi:10.1002/cne.902550407

Brembs, B., Christiansen, F., Pflüger, H.J., Duch, C., 2007. Flight initiation and maintenance deficits in flies with genetically altered biogenic amine levels. J. Neurosci. 27, 11122–11131. doi:10.1523/JNEUROSCI.2704-07.2007

Campbell, H.R., Thompson, K.J., Siegler, M.V., 1995. Neurons of the median neuroblast lineage of the grasshopper: a population study of the efferent DUM neurons. J. Comp. Neurol. 358, 541–551. doi:10.1002/cne.903580407

Charvet, C.J., Striedter, G.F., Finlay, B.L., 2011. Evo-devo and brain scaling: candidate developmental mechanisms for variation and constancy in vertebrate brain evolution. Brain Behav. Evol. 78, 248–257. doi:10.1159/000329851

Chen, C.-L., Hermans, L., Viswanathan, M.C., Fortun, D., Aymanns, F., Unser, M., Cammarato, A., Dickinson, M.H., Ramdya, P., 2018. Imaging neural activity in the ventral nerve cord of behaving adult Drosophila. Nat. Commun. 9, 4390. doi:10.1038/s41467-018-06857-z

Court, R.C., Armstrong, J.D., Borner, J., Card, G., Costa, M., Dickinson, M., Duch, C., Korff, W., Mann, R., Merritt, D., Murphey, R., Namiki, S., Seeds, A., Shepherd, D., Shirangi, T., Simpson, J., Truman, J., Tuthill, J., Williams, D., 2017. A Systematic Nomenclature for the *Drosophila* Ventral Nervous System. BioRxiv. doi:10.1101/122952

Dekkers, M.P.J., Barde, Y.-A., 2013. Developmental biology. Programmed cell death in neuronal development. Science 340, 39–41. doi:10.1126/science.1236152

Doe, C.Q., 1992. Molecular markers for identified neuroblasts and ganglion mother cells in the Drosophila central nervous system. Development 116, 855–863.

Doe, C.Q., Goodman, C.S., 1985. Early events in insect neurogenesis: I. Development and segmental differences in the pattern of neuronal precursor cells. Dev. Biol. 111, 193–205. doi:10.1016/0012-1606(85)90445-2

Doe, C.Q., Kuwada, J.Y., Goodman, C.S., 1985. From epithelium to neuroblasts to neurons: the role of cell interactions and cell lineage during insect neurogenesis. Philos. Trans. R. Soc. Lond. B, Biol. Sci. 312, 67–81. doi:10.1098/rstb.1985.0178

Draizen, T.A., Ewer, J., Robinow, S., 1999. Genetic and hormonal regulation of the death of peptidergic neurons in the Drosophila central nervous system. Dev. Neurobiol.

Duch, C., Pflüger, H.J., 1999. DUM neurons in locust flight: a model system for amine-mediated peripheral adjustments to the requirements of a central motor program. Journal of Comparative Physiology A: Sensory, Neural, and Behavioral Physiology 184, 489–499. doi:10.1007/s003590050349

Dudley, R., 2002. Chapter 6: Evolution of flight and flightlessness, in: The‘ ‘biomechanics of insect flight: Form, function, evolution. Princeton University Press, Princeton, N.J, pp. 261–301.

Eckert, M., Rapus, J., Nürnberger, A., Penzlin, H., 1992. A new specific antibody reveals octopamine-like immunoreactivity in cockroach ventral nerve cord. J. Comp. Neurol. 322, 1–15. doi:10.1002/cne.903220102

Fricker, M., Tolkovsky, A.M., Borutaite, V., Coleman, M., Brown, G.C., 2018. Neuronal Cell Death. Physiol. Rev. 98, 813–880. doi:10.1152/physrev.00011.2017

Greer, C.L., Grygoruk, A., Patton, D.E., Ley, B., Romero-Calderon, R., Chang, H.-Y., Houshyar, R., Bainton, R.J., Diantonio, A., Krantz, D.E., 2005. A splice variant of the Drosophila vesicular monoamine transporter contains a conserved trafficking domain and functions in the storage of dopamine, serotonin, and octopamine. J. Neurobiol. 64, 239–258. doi:10.1002/neu.20146

Hagan, H.R., 1951. Chapter 8: Adenotrophic Viviparity - Diptera (Pupipara), in: Embryology of the Viviparous Insects. The Ronald Press Company, New York, pp. 159–205.

Harris, R.M., Pfeiffer, B.D., Rubin, G.M., Truman, J.W., 2015. Neuron hemilineages provide the functional ground plan for the Drosophila ventral nervous system. Elife 4. doi:10.7554/eLife.04493

Hartenstein, V., Campos-Ortega, J.A., 1984. Early neurogenesis in wild-typeDrosophila melanogaster. Wilhelm Roux’. Archiv. 193, 308–325. doi:10.1007/BF00848159

Herculano-Houzel, S., Manger, P.R., Kaas, J.H., 2014. Brain scaling in mammalian evolution as a consequence of concerted and mosaic changes in numbers of neurons and average neuronal cell size. Front. Neuroanat. 8, 77. doi:10.3389/fnana.2014.00077

Horder, T.J., 1989. Syllabus For An Embryological Synthesis, in: Wake, D.B., Roth, G. (Eds.), Complex Organismal Functions: Integration and Evolution in Vertebrates. Wiley.

Hutson, A.M., 1984. Keds, flat-flies and bat-flies, in: Handbooks for the identification of British insects. Royal Entomological Society, London.

Imms, A.D., 1942. On Braula coeca Nitsch and its affinities. Parasitology 34, 88. doi:10.1017/S0031182000016012

Jia, X.X., Siegler, M.V.S., 2002. Midline lineages in grasshopper produce neuronal siblings with asymmetric expression of Engrailed. Development 129, 5181–5193.

Karcavich, R., Doe, C.Q., 2005. Drosophila neuroblast 7-3 cell lineage: a model system for studying programmed cell death, Notch/Numb signaling, and sequential specification of ganglion mother cell identity. J. Comp. Neurol. 481, 240–251. doi:10.1002/cne.20371

Kimura, K.I., Truman, J.W., 1990. Postmetamorphic cell death in the nervous and muscular systems of Drosophila melanogaster. J. Neurosci. 10, 403–401.

Kondo, S., Senoo-Matsuda, N., Hiromi, Y., Miura, M., 2006. DRONC coordinates cell death and compensatory proliferation. Mol. Cell. Biol. 26, 7258–7268. doi:10.1128/MCB.00183-06

Konings, P.N., Vullings, H.G., Geffard, M., Buijs, R.M., Diederen, J.H., Jansen, W.F., 1988. Immunocytochemical demonstration of octopamine-immunoreactive cells in the nervous system of Locusta migratoria and Schistocerca gregaria. Cell Tissue Res. 251, 371–379. doi:10.1007/BF00215846

Kumar, A., Bello, B., Reichert, H., 2009. Lineage-specific cell death in postembryonic brain development of Drosophila. Development 136, 3433–3442. doi:10.1242/dev.037226

Kutsch, W., Breidbach, O., 1994. Homologous Structures in the Nervous Systems of Arthropoda, in: Advances in Insect Physiology Volume 24, Advances in Insect Physiology. Elsevier, pp. 1–113. doi:10.1016/S0065-2806(08)60082-X

Kuwada, J.Y., Goodman, C.S., 1985. Neuronal determination during embryonic development of the grasshopper nervous system. Dev. Biol. 110, 114–126. doi:10.1016/0012-1606(85)90069-7

Lacin, H., Chen, H.-M., Long, X., Singer, R.H., Lee, T., Truman, J.W., 2019. Neurotransmitter identity is acquired in a lineage-restricted manner in the Drosophila CNS. Elife 8. doi:10.7554/eLife.43701

Lacin, H., Truman, J.W., 2016. Lineage mapping identifies molecular and architectural similarities between the larval and adult Drosophila central nervous system. Elife 5, e13399. doi:10.7554/eLife.13399

Lacin, H., Williamson, W.R., Card, G.M., Skeath, J.B., Truman, J.W., 2020. Unc-4 acts to promote neuronal identity and development of the take-off circuit in the Drosophila CNS. Elife 9. doi:10.7554/eLife.55007

Lacin, H., Zhu, Y., Wilson, B.A., Skeath, J.B., 2014. Transcription factor expression uniquely identifies most postembryonic neuronal lineages in the Drosophila thoracic central nervous system. Development 141, 1011–1021. doi:10.1242/dev.102178

Landgraf, M., Sánchez-Soriano, N., Technau, G.M., Urban, J., Prokop, A., 2003. Charting the Drosophila neuropile: a strategy for the standardised characterisation of genetically amenable neurites. Dev. Biol. 260, 207–225. doi:10.1016/S0012-1606(03)00215-X

Lee, G., Sehgal, R., Wang, Z., Nair, S., Kikuno, K., Chen, C.-H., Hay, B., Park, J.H., 2013. Essential role of grim-led programmed cell death for the establishment of corazonin-producing peptidergic nervous system during embryogenesis and metamorphosis in Drosophila melanogaster. Biol. Open 2, 283–294. doi:10.1242/bio.20133384

Lehane, M.J., 2008. Chapter 9: The blood-sucking insect groups, in: The Biology of Blood-Sucking in Insects. Cambridge University Press, Cambridge, pp. 202–258. doi:10.1017/CCOL0521836085.009

Lin, S., Lai, S.-L., Yu, H.-H., Chihara, T., Luo, L., Lee, T., 2010. Lineage-specific effects of Notch/Numb signaling in post-embryonic development of the Drosophila brain. Development 137, 43–51. doi:10.1242/dev.041699

Marin, E.C., Dry, K.E., Alaimo, D.R., Rudd, K.T., Cillo, A.R., Clenshaw, M.E., Negre, N., White, K.P., Truman, J.W., 2012. Ultrabithorax confers spatial identity in a context-specific manner in the Drosophila postembryonic ventral nervous system. Neural Dev. 7, 31. doi:10.1186/1749-8104-7-31

McAlister, E., 2018. Chapter 9: The parasites, in: The secret life of flies. Natural History Museum, London, pp. 185–211.

Mellert, D.J., Williamson, W.R., Shirangi, T.R., Card, G.M., Truman, J.W., 2016. Genetic and environmental control of neurodevelopmental robustness in drosophila. PLoS One 11, e0155957. doi:10.1371/journal.pone.0155957

Monastirioti, M., Gorczyca, M., Rapus, J., Eckert, M., White, K., Budnik, V., 1995. Octopamine immunoreactivity in the fruit fly Drosophila melanogaster. J. Comp. Neurol. 356, 275–287. doi:10.1002/cne.903560210

Monastirioti, M., Linn, C.E., White, K., 1996. Characterization of Drosophila tyramine beta-hydroxylase gene and isolation of mutant flies lacking octopamine. J. Neurosci. 16, 3900–3911.

Pauls, D., Blechschmidt, C., Frantzmann, F., El Jundi, B., Selcho, M., 2018. A comprehensive anatomical map of the peripheral octopaminergic/tyraminergic system of Drosophila melanogaster. Sci. Rep. 8, 15314. doi:10.1038/s41598-018-33686-3

Petersen, D.S., Kreuter, N., Heepe, L., Büsse, S., Wellbrock, A.H.J., Witte, K., Gorb, S.N., 2018. Holding tight to feathers - structural specializations and attachment properties of the avian ectoparasite Crataerina pallida (Diptera, Hippoboscidae). J. Exp. Biol. 221. doi:10.1242/jeb.179242

Peterson, C., Carney, G.E., Taylor, B.J., White, K., 2002. reaper is required for neuroblast apoptosis during Drosophila development. Development 129, 1467–1476.

Pflüger, H.J., Stevenson, P.A., 2005. Evolutionary aspects of octopaminergic systems with emphasis on arthropods. Arthropod Struct Dev 34, 379–396. doi:10.1016/j.asd.2005.04.004

Prieto-Godino, L.L., Silbering, A.F., Khallaf, M.A., Cruchet, S., Bojkowska, K., Pradervand, S., Hansson, B.S., Knaden, M., Benton, R., 2020. Functional integration of “undead” neurons in the olfactory system. Sci. Adv. 6, eaaz7238. doi:10.1126/sciadv.aaz7238

Rakic, P., 2009. Evolution of the neocortex: a perspective from developmental biology. Nat. Rev. Neurosci. 10, 724–735. doi:10.1038/nrn2719

Ramdya, P., Benton, R., 2010. Evolving olfactory systems on the fly. Trends Genet. 26, 307–316. doi:10.1016/j.tig.2010.04.004

Roeder, T., 2005. Tyramine and octopamine: ruling behavior and metabolism. Annu Rev Entomol 50, 447–477. doi:10.1146/annurev.ento.50.071803.130404

Rogulja-Ortmann, A., Lüer, K., Seibert, J., Rickert, C., Technau, G.M., 2007. Programmed cell death in the embryonic central nervous system of Drosophila melanogaster. Development 134, 105–116. doi:10.1242/dev.02707

Rowell, H.F., 1976. The cells of the insect neurosecretory system: constancy, variability, and the concept of the unique identifiable neuron, in: Advances in insect physiology. Elsevier, pp. 63–123. doi:10.1016/S0065-2806(08)60254-4

Schlurmann, M., Hausen, K., 2003. Mesothoracic ventral unpaired median (mesVUM) neurons in the blowfly Calliphora erythrocephala. J. Comp. Neurol. 467, 435–453. doi:10.1002/cne.10930

Shepherd, D., Bate, C.M., 1990. Spatial and temporal patterns of neurogenesis in the embryo of the locust (Schistocerca gregaria). Development.

Shepherd, D., Harris, R., Williams, D., Truman, J.W., 2016. Postembryonic Lineages of the Drosophila Ventral Nervous System: Neuroglian expression reveals the adult hemilineage associated fiber tracts in the adult thoracic neuromeres. J. Comp. Neurol. 524, 2677–2695. doi:10.1002/cne.23988

Shepherd, D., Sahota, V., Court, R., Williams, D.W., Truman, J.W., 2019. Developmental organization of central neurons in the adult Drosophila ventral nervous system. J. Comp. Neurol. 527, 2573–2598. doi:10.1002/cne.24690

Siegler, M.V., Manley Jr., P.E., Thompson, K.J., 1991. Sulphide silver staining for endogenous heavy metals reveals subsets of dorsal unpaired median (DUM) neurons in insects. J Exp Biol 157, 565–571.

Siegler, M.V., Pankhaniya, R.R., 1997. Engrailed protein is expressed in interneurons but not motor neurons of the dorsal unpaired median group in the adult grasshopper. J. Comp. Neurol. 388, 658–668.

Siegler, M.V., Pankhaniya, R.R., Jia, X.X., 2001. Pattern of expression of engrailed in relation to gamma-aminobutyric acid immunoreactivity in the central nervous system of the adult grasshopper. J. Comp. Neurol. 440, 85–96. doi:10.1002/cne.1371

Sombati, S., Hoyle, G., 1984. Generation of specific behaviors in a locust by local release into neuropil of the natural neuromodulator octopamine. J. Neurobiol. 15, 481–506. doi:10.1002/neu.480150607

Southwell, D.G., Paredes, M.F., Galvao, R.P., Jones, D.L., Froemke, R.C., Sebe, J.Y., Alfaro-Cervello, C., Tang, Y., Garcia-Verdugo, J.M., Rubenstein, J.L., Baraban, S.C., Alvarez-Buylla, A., 2012. Intrinsically determined cell death of developing cortical interneurons. Nature 491, 109–113. doi:10.1038/nature11523

Spörhase-Eichmann, U., Vullings, H.G., Buijs, R.M., Hörner, M., Schürmann, F.W., 1992. Octopamine-immunoreactive neurons in the central nervous system of the cricket, Gryllus bimaculatus. Cell Tissue Res. 268, 287–304.

Stevenson, P.A., Pflüger, H.J., Eckert, M., Rapus, J., 1992. Octopamine immunoreactive cell populations in the locust thoracic-abdominal nervous system. J. Comp. Neurol. 315, 382–397. doi:10.1002/cne.903150403

Stevenson, P.A., Spörhase-Eichmann, U., 1995. Localization of octopaminergic neurones in insects. Comp. Biochem. Physiol. A, Physiol. 110, 203–215. doi:10.1016/0300-9629(94)00152-J

Tamarelle, M., Haget, A., Ressouches, A., 1985. Segregation, division, and early patterning of lateral thoracic neuroblasts in the embryos of Carausius morosus Br. (Phasmida[: Lonchodidae). International Journal of Insect Morphology and Embryology 14, 307–317. doi:10.1016/0020-7322(85)90045-5

Thomas, J.B., Bastiani, M.J., Bate, M., Goodman, C.S., 1984. From grasshopper to Drosophila: a common plan for neuronal development. Nature 310, 203–207. doi:10.1038/310203a0

Thompson, K.J., Siegler, M.V., 1991. Anatomy and physiology of spiking local and intersegmental interneurons in the median neuroblast lineage of the grasshopper. J. Comp. Neurol. 305, 659–675. doi:10.1002/cne.903050409

Thompson, K.J., Siegler, M.V., 1993. Development of segment specificity in identified lineages of the grasshopper CNS. J. Neurosci. 13, 3309–3318.

Truman, J.W., 1990. Metamorphosis of the central nervous system of Drosophila. J. Neurobiol. 21, 1072–1084. doi:10.1002/neu.480210711

Truman, J.W., 1996. Metamorphosis of the Insect Nervous System, in: Gilbert, L., Tata, J., Atkinson, B. (Eds.), Metamorphosis: Postembryonic Reprogramming of Gene Expression in Amphibian and Insect Cells. Academic Press.

Truman, J.W., Ball, E.E., 1998. Patterns of embryonic neurogenesis in a primitive wingless insect, the silverfish, Ctenolepisma longicaudata: comparison with those seen in flying insects. Dev. Genes Evol. 208, 357–368. doi:10.1007/s004270050192

Truman, J.W., Bate, M., 1988. Spatial and temporal patterns of neurogenesis in the central nervous system of Drosophila melanogaster. Dev. Biol. 125, 145–157. doi:10.1016/0012-1606(88)90067-X

Truman, J.W., Moats, W., Altman, J., Marin, E.C., Williams, D.W., 2010. Role of Notch signaling in establishing the hemilineages of secondary neurons in Drosophila melanogaster. Development 137, 53–61. doi:10.1242/dev.041749

Truman, J.W., Schuppe, H., Shepherd, D., Williams, D.W., 2004. Developmental architecture of adult-specific lineages in the ventral CNS of Drosophila. Development 131, 5167–5184. doi:10.1242/dev.01371

Truman, J.W., Talbot, W.S., Fahrbach, S.E., Hogness, D.S., 1994. Ecdysone receptor expression in the CNS correlates with stage-specific responses to ecdysteroids during Drosophila and Manduca development. Development 120, 219–234.

Walker, M D, Rotherham, I.D., 2010a. Characteristics of Crataerina pallida (Diptera: Hippoboscidae) populations; a nest ectoparasite of the common swift, Apus apus (Aves: Apodidae). Exp Parasitol 126, 451–455. doi:10.1016/j.exppara.2010.05.019

Walker, Mark D, Rotherham, I.D., 2010b. The common swift louse fly, Crataerina pallida: an ideal species for studying host-parasite interactions. J. Insect Sci. 10, 193. doi:10.1673/031.010.19301

Wheeler, W.M., 1891. Neuroblasts in the arthropod embryo. J Morphol 4, 337–343. doi:10.1002/jmor.1050040305

White, K., Grether, M.E., Abrams, J.M., Young, L., Farrell, K., Steller, H., 1994. Genetic control of programmed cell death in Drosophila. Science 264, 677–683. doi:10.1126/science.8171319

Witten, J.L., Truman, J.W., 1998. Distribution of GABA-like immunoreactive neurons in insects suggests lineage homology. J. Comp. Neurol. 398, 515–528. doi:10.1002/(SICI)1096-9861(19980907)398:4<515::AID-CNE4>3.0.CO;2-5

Yellman, C., Tao, H., He, B., Hirsh, J., 1997. Conserved and sexually dimorphic behavioral responses to biogenic amines in decapitated Drosophila. Proc. Natl. Acad. Sci. USA 94, 4131–4136. doi:10.1073/pnas.94.8.4131

